# Ryanodine receptor 1 (*RYR1*) patient-derived muscle cells recapitulate disease phenotypes in 2D and 3D culture models

**DOI:** 10.64898/2026.06.18.732485

**Authors:** Joshua S. Clayton, Jordan J. Crane, Jeremy D. D. Garcia, Divya Avnoor, Anna C. Johnstone, Ria Aggarwal, Changho Chun, Vanessa Crossman, Peter J. Houweling, Edoardo Malfatti, Norma B. Romero, David L. Mack, Nigel G. Laing, Gianina Ravenscroft, Rhonda L. Taylor

## Abstract

The Ryanodine Receptor 1 (RyR1) is the major Ca^2+^ release channel in skeletal muscle and plays a crucial role in excitation-contraction coupling. Pathogenic variants in *RYR1* are the most common cause of congenital myopathy, for which there are no approved treatments. Patient-centric disease models may help to facilitate the design and screening of novel therapeutics in a human genomic context. In this report, we describe the differentiation of five dominant *RYR1*-related myopathy patient-derived induced pluripotent stem cell (iPSC) lines into muscle progenitor cells (MPCs), and subsequently into multinucleated myotubes in 2- and 3- Dimensional (D) culture models. In 2D, we show significantly reduced Ca^2+^ release in a patient line compared to a healthy control following stimulation with caffeine. In 3D engineered muscle tissues (EMTs), patient-relevant phenotypes including reduced twitch amplitude, delayed relaxation and altered force-frequency relationships were observed in a patient line compared to two healthy controls. We also show that the 2D cultures are a suitable platform for screening the efficacy and cellular toxicity of antisense oligonucleotide therapeutics. Together, these results suggest that iPSC-derived skeletal muscle cultures are useful models for understanding the pathobiology of *RYR1*-related myopathies and as a testbed for emerging treatments.

## Introduction

The Ryanodine Receptors (RyRs) are a family of homotetrameric Ca^2+^ release channels, that are located on the sarcoplasmic/endoplasmic reticulum (SR/ER) membrane where they function to control the release of Ca^2+^ from intracellular stores into the cytosol^1^. In mammals, there are three RyR isoforms with distinct expression patterns; RyR1 is predominantly expressed in skeletal muscle, RyR2 in cardiac muscle (and to a lesser extent in brain and smooth muscle), and RyR3 is lowly expressed in a range of tissues including skeletal muscle, brain and smooth muscle^1,2^. In skeletal muscle, the RyR1 isoform interacts with the voltage-sensing dihydropyridine receptor (DHPR, also known as Cav1.1) in a dynamic relationship that is central to excitation-contraction coupling, and therefore skeletal muscle function^3–5^. Genetic variants which perturb the function of RyR1, disrupt the synchronous relationship between Ca^2+^ signalling, muscle contraction and muscle relaxation.

Dominant *RYR1* variants are the leading cause of malignant hyperthermia susceptibility (MHS; MIM#145600), a potentially fatal pharmacogenetic disorder triggered by some anaesthetics^6,7^ with an estimated population prevalence of ∼1/2,000 - 1/80,000^8,9^. MHS individuals may be completely asymptomatic until exposed to a triggering agent, or more rarely, may have a concurrent congenital myopathy^10,11^. Pathogenic variants in *RYR1* are also the most common cause of congenital myopathy, affecting approximately 1/90,000 people worldwide^12–14^. Historically, diagnoses were guided by histological features on muscle biopsy including central core disease (CCD; MIM#117000)^15^, multi-minicore disease (MmD; MIM#255320)^11,16^, dusty core disease (DuCD)^17^, centronuclear myopathy^18^ and others^19,20^. However, an ever-expanding association with overlapping pathologies and phenotypes such as King-Denborough Syndrome (MIM#619542) and fetal akinesias^21,22^ has led to the term *RYR1*-related myopathies (*RYR1*-RM) being used to collectively describe the wide-spectrum of congenital muscle diseases associated with *RYR1* pathology^23–25^. Together, *RYR1*-RM are a heterogeneous group of skeletal muscle diseases, primarily characterised by slowly- or non-progressive muscle weakness^10,11,26^. The spectrum of clinical features is diverse and can include fetal hypokinesia, delayed motor milestones, muscle cramps, myalgia, joint contractures, scoliosis, facial weakness, ophthalmoparesis, bulbar involvement and respiratory insufficiency^10,11^.

It is thought that *RYR1*-RM may be caused by at least four different mechanisms; 1) uncontrolled leak of Ca^2+^ through RyR1^27^, 2) increased channel sensitivity to activators^27–29^, 3) excitation-contraction uncoupling leading to reduced Ca^2+^ release through RyR1^30^, and 4) reduced RyR1 abundance^31,32^. Ultimately, each of these mechanisms dysregulates cytosolic Ca^2+^ homeostasis leading to reduced muscle contractile performance in response to an action potential. Therefore, considerable effort has been applied to develop treatments that normalise RyR1 channel function.

There are currently no disease-modifying treatments available for *RYR1*-RM, although a recent phase I open label trial of Rycal S48168 (ARM210) was well tolerated and improved some key disease features (fatigue, proximal muscle weakness) in four patients treated with the highest dose^33^. Preclinically, allopurinol and xanthine derivatives have been investigated as potential therapeutics that might stabilise and enhance activity of RyRs in the context of loss-of-function or reduced-expression variants^34^. RNA modulating therapies that selectively target specific patient variants, CRISPR-based pathogenic variant correction by prime-editing, and a variant-agnostic approach to selectively remove pathogenic *RYR1* gene copies using CRISPR/Cas9 are also under investigation^35–38^. However, there remains a need for the continued development of targeted treatments, as well as improved models for pre-clinical screening of novel treatments.

*RYR1*-RM disease mechanisms have previously been studied using 1) transient delivery of *RYR1* cDNA into cultured cells (HEK293, or RyR1-null murine myotubes), 2) RyR1-mediated Ca^2+^ transients in non-muscle cells (lymphoblastoid cells expressing low levels of RyR1), and 3) isolation and culture of human primary cells from muscle biopsy^39,40^. There are also numerous RyR1 mouse models encompassing heterozygous, compound heterozygous and knockout variants^40^. While these models have undoubtedly been useful in generating biological insights, there is considerable scope for development of patient-centric models that reflect the human genomic context while still accurately modelling the primary outcome measures of *RYR1*-RM such as muscle weakness. To achieve this, many researchers are now using patient-derived induced pluripotent stem cells (iPSCs) that provide a near limitless source of patient cells^41^. Several groups have established methods for differentiation of iPSCs to skeletal muscle^42–44^ and iPSC-derived skeletal muscle cells are already in use to model other muscle diseases and test treatments^45–48^. We reasoned that patient iPSC-derived myotubes may be a useful model to study *RYR1*-RM and test potential therapies.

## Results

### Assessment of RYR1 variants in a cohort of five patients diagnosed with CCD with or without concurrent MHS

We have previously obtained lymphoblastoid cell lines (LCLs) from five unrelated *RYR1*-RM patients (Généthon, France) and reprogrammed these to generate iPSC lines^49–51^. All five patients received a clinical diagnosis of CCD, and two also had concurrent MHS (Table 1). In all five cases, genetic testing identified heterozygous variants in *RYR1*. Of these five variants, two are known recurrent missense variants (p.V2168M^28,52,53^ and p.R2508C^15,54^), one has previously been reported as a *de novo* variant (p.H4813Y^55^), and the remaining two (p.N1346K and p.N4715-18del) have not been previously described as pathogenic (outside of our own iPSC publications).

**Table 1.**
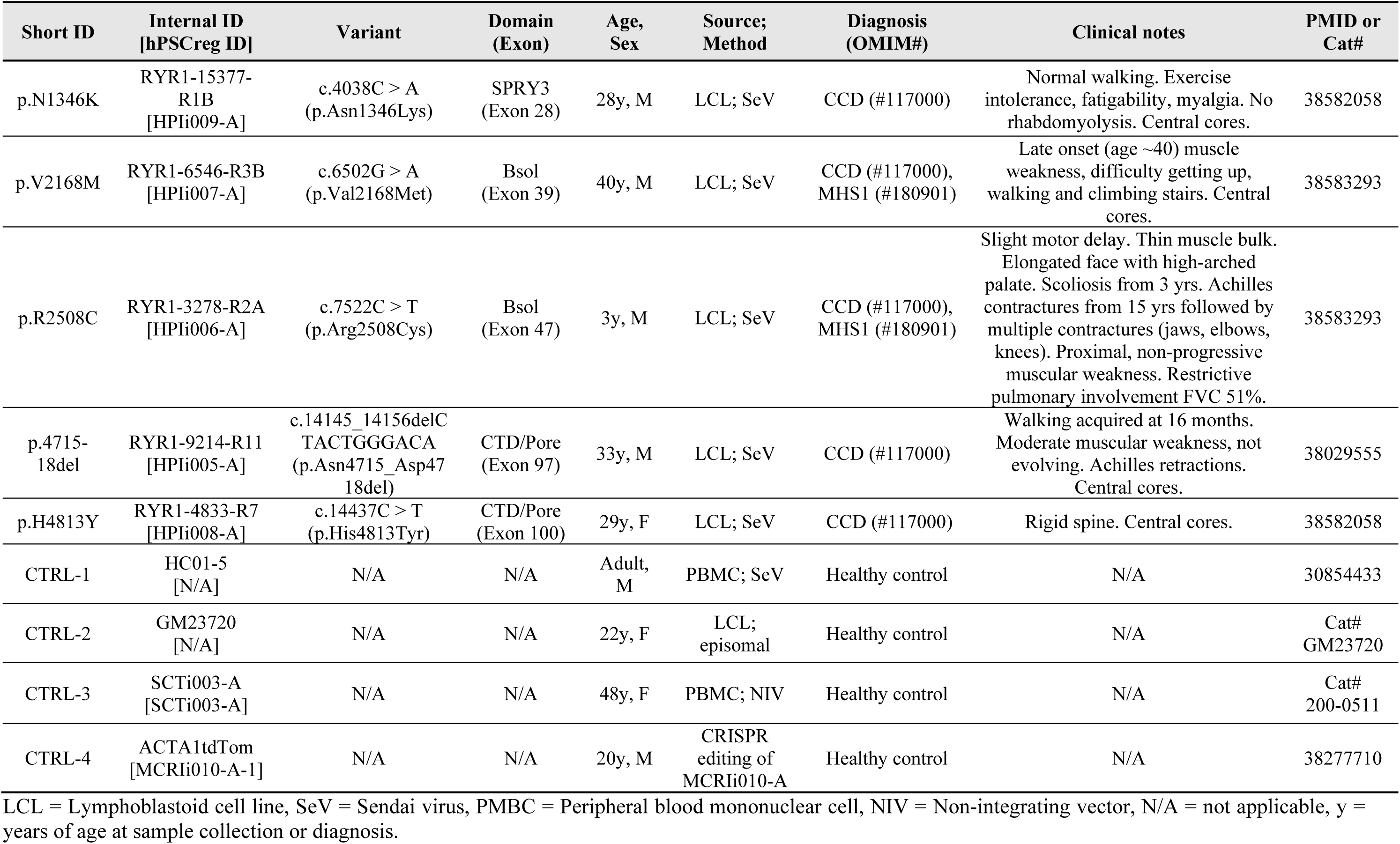
Patient and cell line information.

Structurally, the p.N1346K substitution sits in the Divergent Region 2 (D2) within the SPRY3 domain^56^ (Fig. 1A), which is implicated in tetrad formation^57^. Both the p.N4715-18del and the p.H4813K substitution are located in the transmembrane portion of RyR1 (Fig. 1A, B; Note that the p.N1346 amino acid is in an area of the protein not modelled in the crystal structure so is not visualised). The effect/s of these three protein changes (p.N1346K, p.N4715-18del, p.H4813K) on channel function have not yet been studied.

**Figure 1.**
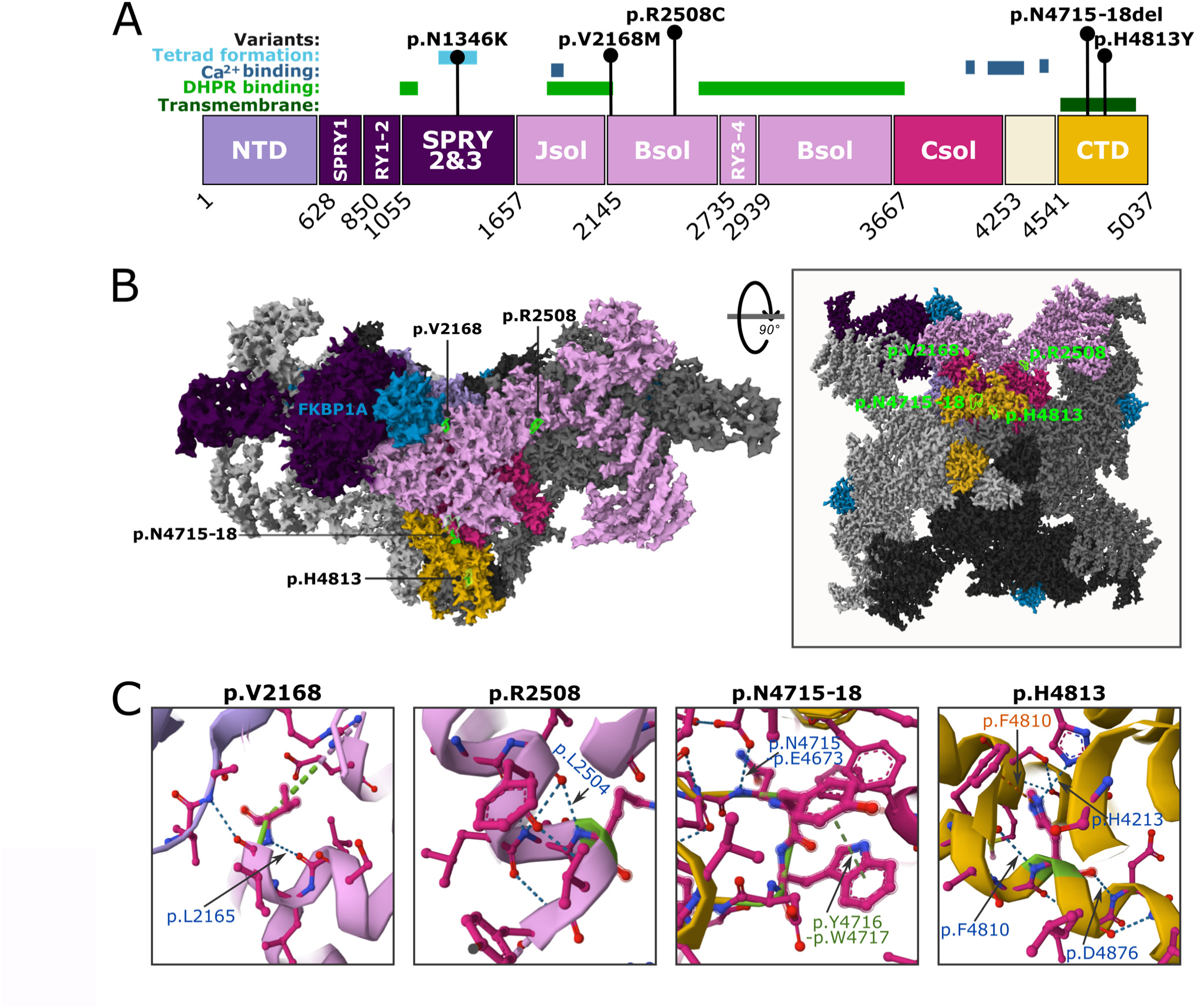
*RYR1*-RM patient variants mapped onto the RyR1 protein structure. **A**. Linear schematic of the RyR1 protein domains, with key interaction sites and variants annotated. Patient variants are described using Human Genome Variation Society (HGVS) nomenclature. NTD = N-terminal domain, SPRY = SP1a/ryanodine receptor domains, RY = RyR repeat pairs, Jsol = junctional solenoid, Bsol = bridge solenoid, Csol = core solenoid, CTD = C-terminal domain. Schematic modified from others^57^. **B**. Patient variants (highlighted in green) mapped on to the crystal structure of rabbit RyR1 in complex with FKBP12 (blue) at 3.8 Å (PDB ID: 3J8H^80^). A single RyR1 monomer is colourised and annotated, the remaining three monomers are monochromatic. **C**. Bonding interactions of modelled variants are indicated. Blue text indicates a hydrogen bond, green text indicates a Pi stacking interaction, orange text indicates cation-Pi-stacking interaction. The p.N1346K substitution sits within an unmodelled part of the crystal structure and is not shown.

The p.V2168M substitution is situated within the bridge solenoid domain (Bsol, Fig. 1A, B), in a region predicted to interact with DHPR^57^. This variant is commonly associated with MHS without CCD^28,52^, although one concurrent CCD and MHS case has been reported where the patient presented with proximal weakness, mild orthopaedic issues during infancy and cores in type I myofibres^53^. The p.R2508C substitution is also located in the Bsol domain (Fig. 1A, B) and is known to cause CCD concurrent with MHS, with phenotypes including proximal muscle weakness, scoliosis, contractures (Table 1), congenital arthrogryposis, and cores with type I myofibre predominance on muscle biopsy^54^.

All modelled variants were shown to participate in hydrogen bonding with some additionally involving Pi stacking or cation-Pi stacking interactions (Fig. 1C). This is in line with the observations of Chang *et al*.^58^ who found that dominant *RYR1* variants were most likely to act *via* disruption of hydrogen bonding.

### RYR1 patient-derived iPSC lines can be differentiated to myogenic progenitor cells and multinucleated myotubes in 2D culture

To create patient-derived and control myogenic progenitor cells (MPCs), iPSCs from all five unrelated RYR1-RM patients^49–51^ and three healthy controls (iPSC-CTRL-1, iPSC-CTRL-2, iPSC-CTRL-3) were differentiated using the STEMdiff^TM^ myogenic progenitor kit. After 30 days, single cells representing a mixed population of mononuclear cells were isolated (Fig 2A, Supp. Fig. S1). Patient-derived mononuclear cells were generally similar in morphology to three independent healthy control lines and had a comparable or greater proportion of proliferative (Ki67-positive) cells (Fig. 2B, Supp. Fig S2, Supp. Table S1). Cultures from patients and controls had a low and variable proportion of desmin-positive cells (Fig. 2B, Supp. Fig S2, Supp. Table S1). Given that desmin expression increases during human skeletal muscle lineage progression^59^ we considered that most of these progenitor cells may be developmentally ‘earlier’ in the myogenic trajectory (prior to induction of desmin and MyoD expression) and may therefore express muscle stem cell markers such as SIX1 and/or PAX3^60^.

**Figure 2.**
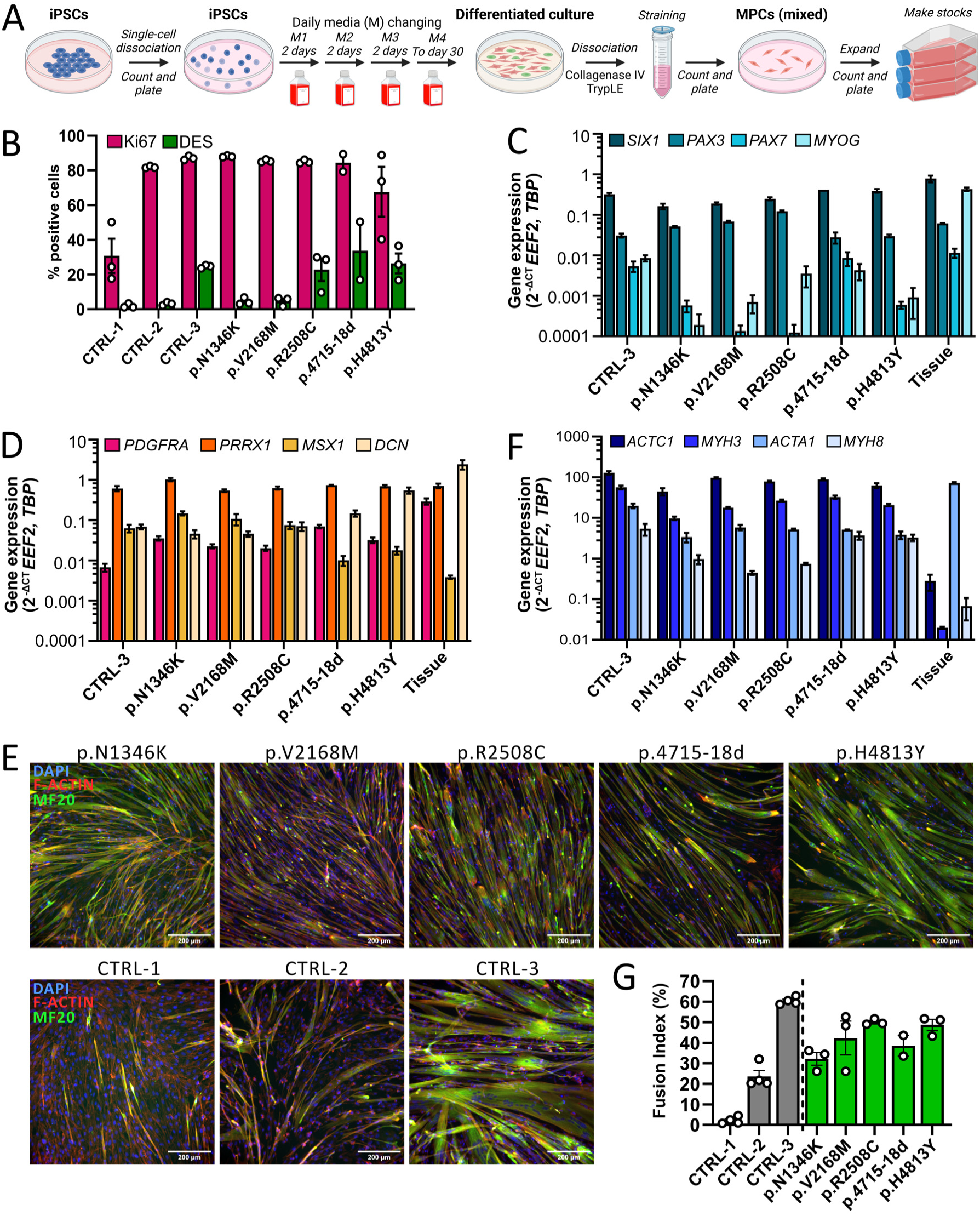
*RYR1*-RM patient derived MPCs differentiate into multinucleated myotubes. **A**. Schematic of the iPSC-to-skeletal muscle differentiation workflow using the STEMdiff myogenic progenitor kit. MPC = myogenic progenitor cell. **B.** Percentage of cells staining positively for a cell proliferation marker (Ki67, pink) and myoblast marker (desmin/DES, green) from microscope images (minimum of 20 images per 96-well). **C**/**D/F**. Relative expression of various transcripts was determined using qRT-PCR. Transcript abundance was calculated relative to the geometric mean of two endogenous control genes (*EEF2* and *TBP*) and is plotted alongside data generated from three independent adult skeletal muscle tissue samples (tissue, *vastus lateralis*). **E**. Day 4 myotubes were fixed and stained for F-actin (phalloidin, red), pan-myosin (MF-20, green) and nuclei (DAPI, blue). A representative image is shown for each line. Scale bars are 200 µm. **G**. Percentage of nuclei overlapping the myosin stain (fusion index; minimum of 20 images per 96-well). Data are mean±SEM for *n* = 3 independent replicates/line, except p.4715-18del which is *n* = 2.

We analysed the abundance of myogenic transcription factors (*SIX1*, *PAX3*, *PAX7*, *MYOG*) by qRT-PCR in all five RYR1-RM patient derived mononuclear cells and compared expression levels to healthy control derived cells (CTRL-3 only) and primary adult skeletal muscle tissues (*vastus lateralis* muscle from n=3 unrelated donors). *SIX1* and *PAX3* were the most abundant myogenic transcription factors in control and patient derived cells (Fig. 2C). Expression of *PAX7* and *MYOG* were comparably lower and more variable between the cell lines (Fig. 2C) supporting our hypothesis that the generated MPCs are likely to be developmentally early progenitors. We also analysed expression of additional markers representative of alternative cell lineages and progenitors expected to be present^44,61^. These included mesenchymal stem cells (*PDGFRA, PRRX1*^62^), progenitors of the neural crest lineage (*MSX1*^63^), and fibroblasts (*DCN*^64^) (Fig. 2D). Of these, expression of *PRRX1,* a marker of somitic mesenchyme in the early limb bud^65^, was most abundant across all samples (Fig. 2D).

To examine their myogenic potential, MPCs were grown to >80% confluence and subsequently cultured for up to 7 days in a low serum media to induce terminal differentiation. By day (D) 4, all lines demonstrated fusion of MPCs into multinucleated myotubes (Supp. Fig. S3) staining positively for myosin heavy chain (Fig. 2E). Between D4 and D7, myotubes of some cultures began to spontaneously lift off the culture plate (Supp. Fig. S4). Thus, quantitative analyses are limited to the D4 timepoint, which also coincides with peak *RYR1* expression reported by others^66^. As a measure of the relative maturity of the myotubes, we used qRT-PCR to quantify the transcript abundance of sarcomeric protein isoforms predominant at fetal/embryonic/neonatal (*ACTC1*, *MYH3, MYH8*) *versus* adult (*ACTA1*) stages of development (Fig. 2F). The expression of the fetal sarcomeric actin isoform, *ACTC1*^67^, was ∼5-20 fold higher than the adult skeletal muscle actin isoform, *ACTA1* (Fig. 2F). Similarly, the expression of the embryonic myosin *MYH3* was higher than the fetal/neonatal myosin *MYH8*^68^ in all lines (Fig. 2F). In accordance with the findings of others^61^, and compared to the expression profile in the adult tissues (Fig. 2F), these data suggest that cultured myotubes in 2D have an embryonic-like level of maturity. Average fusion index for the *RYR1*-patient derived lines ranged from 32-50% (Fig. 2G, Supp. Table S1), which is within the expected range^43^. Together, these analyses confirm the myogenic capacity and quality of our iPSC-derived MPCs.

### RyR1 p.R2508C myotubes produced a Ca^2+^ transient following stimulation with caffeine

We next sought to determine whether our newly created muscle cell lines are a suitable model of *RYR1*-RM. We investigated whether *RYR1* was the predominant ryanodine receptor isoform expressed in our patient-derived myotubes (compared to *RYR3* which is also known to be expressed in skeletal muscle^1^), and whether it was co-expressed with the dihydropyridine receptor (DHPR). We used qPCR to quantify the transcript abundance of *RYR1*, *RYR3* and *CACNA1S* (the alpha1S subunit of the DHPR) (Fig. 3A). In each line, *RYR1* was the predominant ryanodine receptor, consistently expressed at levels ∼50-100 times higher than *RYR3,* and in accordance with the relative abundance of these two transcripts in adult skeletal muscle tissue (Fig. 3A). In all lines, expression of *CACNA1S* was also readily detected (Fig. 3A).

**Figure 3.**
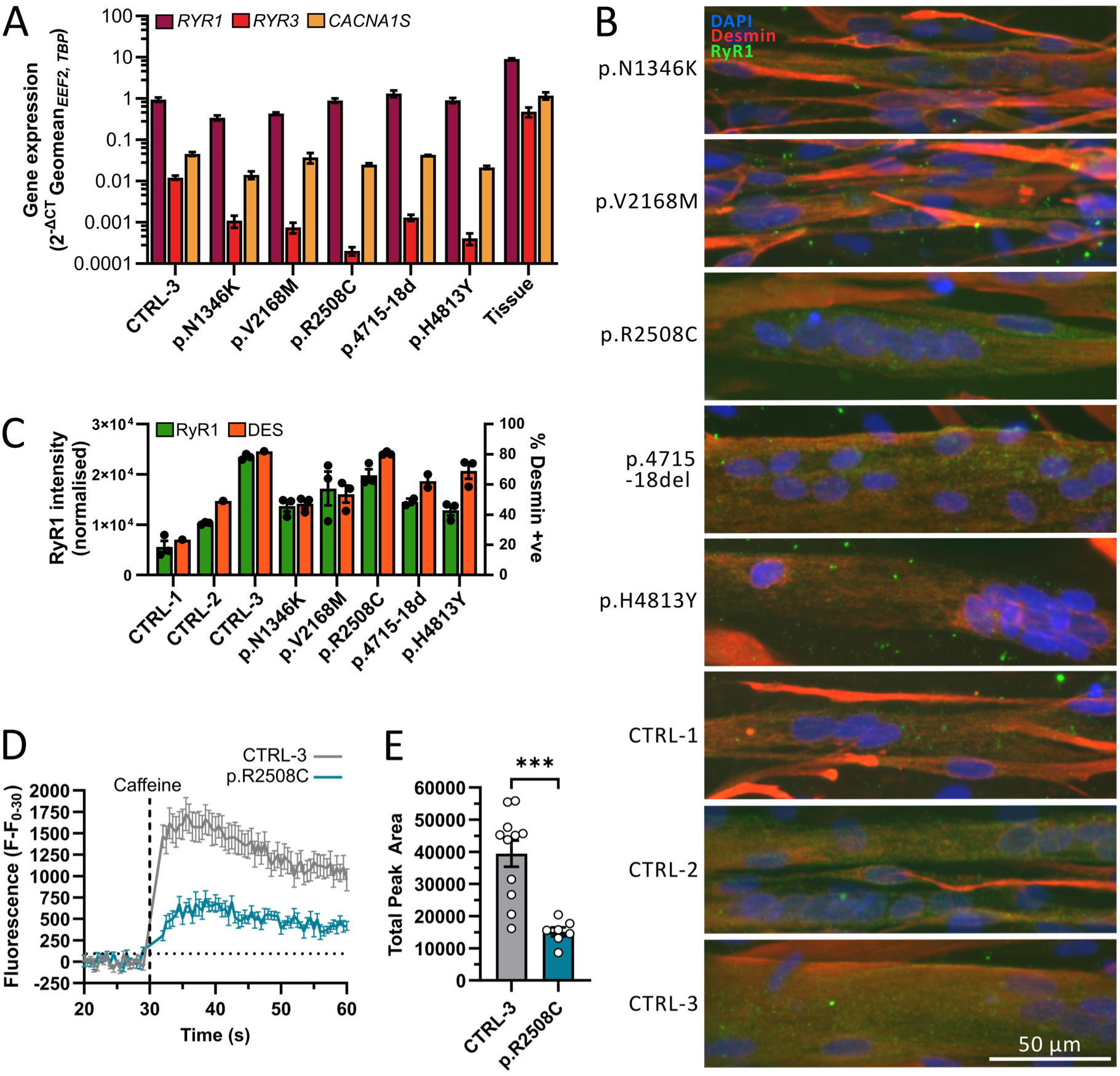
*RYR1*-myotubes harbouring the p.R2508C substitution generate reduced Ca^2+^ release following stimulation with caffeine. **A.** Relative expression of *RYR1*, *RYR3* and *CACNA1S* transcripts were determined using qRT-PCR. Transcript abundance was calculated relative to the geometric mean of two endogenous control genes (*EEF2* and *TBP*) and plotted alongside data generated from three independent adult skeletal muscle tissue samples (tissue, *vastus lateralis*). Data are mean±SEM for n=3 independent replicates/line, except p.4715-18del which is n=2. **B**. Day 4 myotubes were fixed and stained for desmin (red), RyR1 (34C, green) and nuclei (DAPI, blue). Scale bars are 50 µm and apply to all images. **C**. CellProfiler was used to quantify the percentage of nuclei falling within desmin-positive areas of the culture (orange, right y axis), and the relative RyR1 staining intensity within desmin-positive areas (green, left y axis). A minimum of 20 images per 96-well were used for quantification. Data are mean±SEM for n=3 independent replicates/line, except p.4715-18del which is n=2. **D-E**. Myotubes were cultured on hydrogels in 96-well plates and at D14-20 were incubated with Fluo-4 prior to stimulation with 20mM of caffeine. **D**. Ca^2+^ response (Fluo-4 signal) in the whole well over 60 seconds was measured, and **E**. total area under the curve was calculated. Data are mean±SEM of *n* = 11 (CTRL-3) or *n* = 7 wells (p.R2508C) from at least *n* = 2 independent experiments. Statistical comparisons are unpaired t-test with significance set at *p* < 0.05. Welch’s correction was performed when the variance between groups was significantly different.

To determine whether expression of *RYR1* transcript was translated to detectable levels of RyR1 protein, D4 myotubes were co-stained with RyR1 antibody (34C) and desmin (Fig. 3B, C). Positive RyR1 staining could be observed throughout myotubes of all cultures (Fig. 3B, Supp. Fig. S5). Qualitatively, punctate RyR1 staining was localised peripherally and at the myotube surface, as has been reported previously^36,69^. A custom CellProfiler pipeline was used to quantify the intensity of RyR1 staining, as a measure of RyR1 protein abundance in desmin-positive myotubes (Fig. 3C).

Of our five *RYR1* patient derived lines, the p.R2508C variant is the most extensively studied and has been shown to respond to stimulation with caffeine in other model systems^28,29^. Thus we used the p.R2508C line as an exemplar to explore potential RyR1 phenotypes in 2D using a whole-well Ca^2+^ release assay. We selected CTRL-3 as the most suitable control line based on comparable fusion index and RyR1 abundance relative to p.R2508C patient-derived myotubes (Fig. 2G, 3C). Differentiated myotubes were incubated with Fluo-4, a fluorescent Ca^2+^ indicator^70^. The Fluo-4 signal for each well was recorded before, during and after the transient response to stimulation with caffeine (20 mM), an RyR1 agonist.

To prevent myotube loss, we cultured myotubes on a hydrogel which also had the benefit of enabling Ca^2+^ assays to be performed at later time-points when myotubes were more mature^71^. At D14 - D20 of differentiation, we tested the ability of p.R2508C myotubes to produce Ca^2+^ transients in response to caffeine (Fig. 3D). Under these conditions, we found that healthy control myotubes were more likely to respond; for CTRL-3, 92% of wells recorded a positive Ca^2+^ transient (11/12 wells across two independent experiments), compared to only 58% of wells from the p.R2508C line (7/12 wells across two independent experiments). For the wells that produced a Ca^2+^ transient, area under the curve was used as a measure of total Ca^2+^ release^38^. CTRL-3 myotubes released significantly more Ca^2+^ than p.R2508C myotubes (Fig. 3E; Unpaired t-test with Welch’s correction, *p =* 0.0001).

Collectively, we show that *RYR1*-RM patient MPCs can differentiate into myotubes that express RyR1 and are capable of producing Ca^2+^ transients following stimulation with a known RyR1 agonist. Further, reduced total Ca^2+^ release in the p.R2508C myotubes compared to CTRL-3 myotubes suggests that quantifiable deficits in RyR1 channel function can be measured in 2D cultures.

### RYR1 p.R2508C patient-derived 3D EMTs have reduced force output and altered relaxation dynamics

To further assess the utility of our *RYR1*-RM lines to model disease, we generated 3D engineered muscle tissues (EMTs) using MPCs from p.R2508C and two controls; CTRL-3, and an ACTA1-tdTomato reporter line (CTRL-4; Table 1)^72^. CTRL-4 MPCs were generated at a separate site using identical culture and differentiation protocols and were validated to have comparable fusion index (54.9%) in 2D to the p.R2508C line (Supp. Fig. S6). 3D EMTs were generated using the Mantarray^TM^ instrument and cultured for 21 days with longitudinal assessment of contractile function.

Compared to controls, p.R2508C EMTs had ∼70% lower maximum and mean twitch force across all timepoints from D10 to D21 (Fig 4A, 4B; Tables S2, S3). At endpoint (D21), p.R2508C tissues had significantly lower mean twitch force (133.4 ± 8.1 μN, *n* = 4) compared with CTRL-3 (508.3 ± 14.4 μN, *n* = 4, p < 0.0001) and CTRL-4 (478.8 ± 28.7 μN, *n* = 3, *p* = 0.008). Maximum force was similarly lower (195.9 ± 12.2 μN) than CTRL-3 (691.4 ± 14.7 μN, *p* < 0.0001) and CTRL-4 (712.1 ± 52.9 μN, *p* = 0.0146). Controls did not significantly differ at this timepoint.

**Figure 4.**
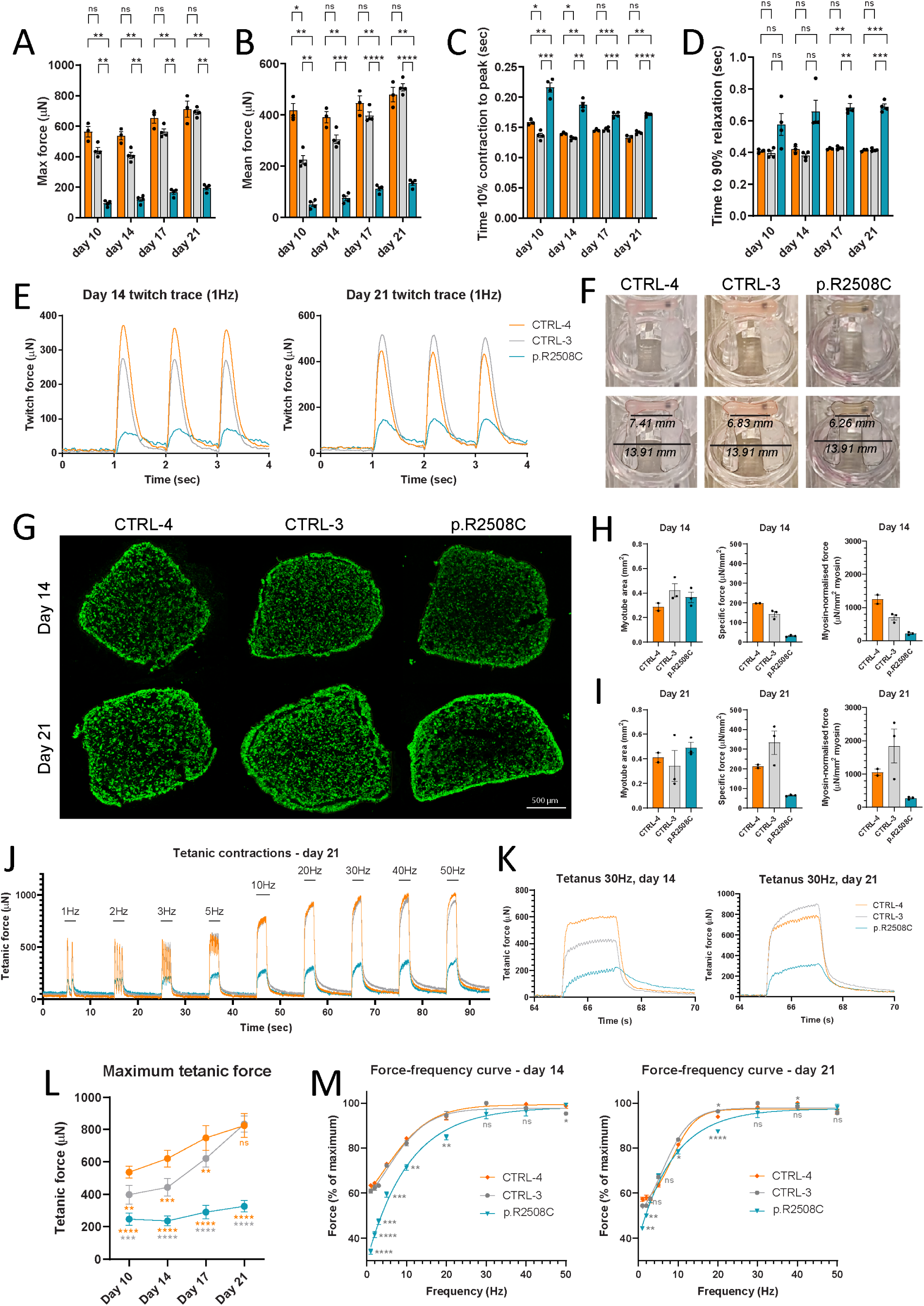
Characterisation of *RYR1* 3D engineered muscle tissues show deficits in force output and relaxation. 3D EMT data for CTRL-4 (ACTA1-tdTomato) - orange, CTRL-3 - grey, and p.R2508C - teal. All data are longitudinal i.e. stimulation data for the same set of EMTs at each timepoint, from *n* = 4 (CTRL-3, p.R2508C) or *n* = 3 (CTRL-4) tissues. **A-D)** Longitudinal twitch data (1Hz stimulation) for EMTs at day 10, 14, 17 and 21 showing maximum twitch **(A)**, mean force **(B)**, time from 10% of contraction to peak; labelled as time to peak for simplicity **(C)** and time from peak to 90% relaxation **(D)**. **E)** Representative twitch traces at day 14 (left) and day 21 (right). **F)** Photos of representative EMTs at day 14. RYR1 p.R2508C tissues show passive stiffness – in lower images EMTs and posts are outlined in black for ease of visualisation, and tissue stiffness is indicated by scaled measurement of the distance between the two Mantarray posts (scaled relative to fixed well diameter of 13.91mm). **G)** Representative cross-sections of 3D EMTs stained with MF20 (pan-myosin; green). Hoechst (nuclei) staining not shown for clarity. Scale bar applies to all EMTs. **H/I)** Normalised force data for day 14 **(H)** and day 21 **(I)** tissues. From left to right: myotube area (MF20-positive as fraction of total area), specific force (mean twitch force normalised to total cross-sectional area), and myosin-normalised force (mean twitch force normalised to myotube (myosin-positive) area). Data shown are for representative EMTs at each timepoint; *n* = 1 tissue, *n* = 2–3 sections. Stats are not included as data only represent *n* = 1 tissue at each timepoint. **J)** Representative tetanus traces at day 21. Stimulation frequencies are noted above each set of corresponding peaks. **K)** Representative 30Hz tetanus traces at day 14 (left) and day 21 (right). **L)** Maximum tetanic force (1-50Hz stimulation window) at day 10, 14, 17 and 21. Significance values (two-way ANOVA) are indicated below datapoints; the colour indicates which sample set has been compared to (orange: compared to CTRL-4, grey: compared to CTRL3). **M)** Force-frequency curves (1-50Hz) at day 14 (left) and day 21 (right). Data are plotted as % of maximum force to normalise for differences in absolute force. Significance values (two-way ANOVA) compared to CTRL-3 are indicated below or above datapoints for p.R2508C and CTRL-4, respectively. **Statistics:** Statistical comparisons were performed using two-way ANOVA with Tukey’s multiple comparisons test (* *p* <0.05, ** *p* < 0.01, *** *p* < 0.001, **** *p* < 0.0001, ns = not significant).

Contraction and relaxation dynamics were also altered: p.R2508C tissues had significantly increased time to peak (Fig. 4C) and time to 90% relaxation (Fig. 4D) across multiple timepoints (Tables S2, S3). At D21, p.R2508C time from 10% contraction to peak (0.171 ± 0.001 sec) was 21% longer than CTRL-3 (0.141 ± 0.001 sec, p < 0.0001) and 29% longer than CTRL-4 (0.133 ± 0.003 sec, p = 0.0031). Similarly, p.R2508C time to 90% relaxation (0.688 ± 0.017 sec) was 66% slower than CTRL-3 (0.416 ± 0.004 sec, p = 0.0006) and 67% slower than CTRL-4 (0.413 ± 0.004 sec, p = 0.0007). The relaxation deficit was pronounced even on visual examination of twitch traces and was maintained as tissues continued to develop and increase in strength (Fig. 4E). At the gross level, p.R2508C EMTs also had distinct passive tension compared to matched controls which, in contrast, rapidly returned to a relaxed state post-stimulation (Fig. 4F).

To confirm that the lower force output and altered biomechanical properties of p.R2508C tissues were not due to substantially lower myotube fusion and tissue development, we harvested representative tissues at D14 and D21 and stained cross-sections using an anti-myosin heavy chain antibody (MF20). Tissue architecture and myogenic population/fusion did not differ (Fig. 4G): CTRL-3 had the highest proportion of myosin-positive area at D14 (CTRL-3 20.7%, *n* = 3 sections), followed by CTRL-4 (16.0%, *n* = 2 sections) and p.R2508C (14.7%, *n* = 3 sections). At D21, all EMTs were 20-25% myosin-positive by area (CTRL-3 19.7%, *n* = 3; CTRL-4 20.2%, *n* = 2; p.R2508C 23.4%, *n* = 3 sections). Subsequent normalisation of twitch force to effective cross-sectional area (specific force) and myosin area for matched tissues showed p.R2508C was still at least 50% weaker than controls at both timepoints (Fig. 4H, 4I).

To assess tetanus and force-frequency relationships, we stimulated tissues with increasing frequency (1-50 Hz) (Fig. 4J). Similar to twitch, p.R2508C tissues had lower force amplitudes and were much slower to relax – as exemplified by stimulation traces; particularly noticeable at D14 (Fig. 4K). Maximum tetanic force was 2- to 4-fold lower compared to controls across all timepoints (Fig. 5L). Force of D21 p.R2508C tissues (326.0 ± 17.8 μN) was 39% of CTRL-3 (834.0 ± 23.3 μN, *p* < 0.0001) and 40% of CTRL-4 (824.4 ± 43.1 μN, *p* < 0.0001). Although force of p.R2508C tissues increased from D10 to D21 (32%), this was much less pronounced compared to CTRL-3 (109% increase) and CTRL-4 (54% increase).

**Figure 5.**
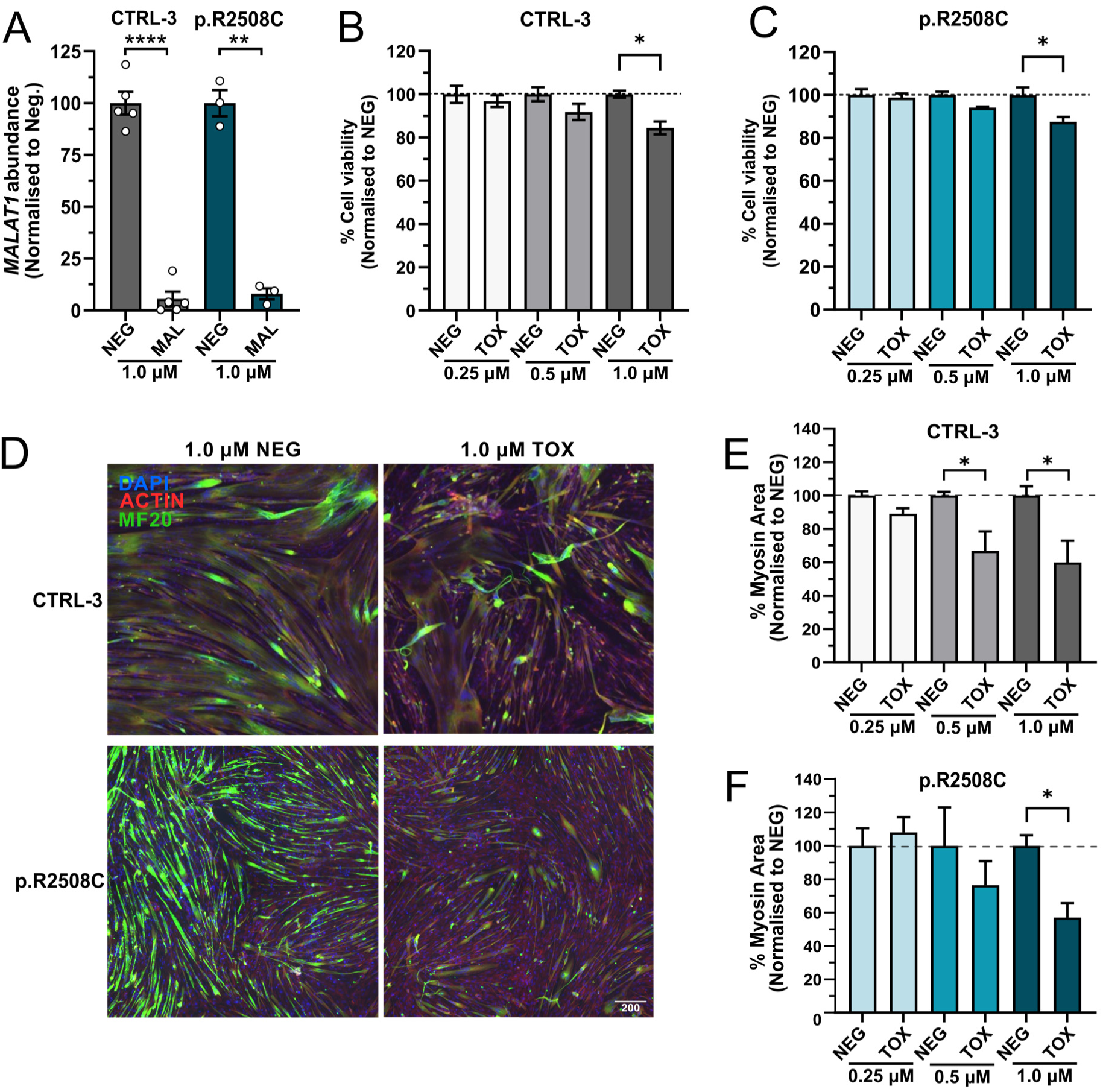
*RYR1*-RM myotubes readily take up ASOs and are amenable to high-throughput screening of knockdown and cell viability. *MALAT1*-FAM^+^ (MAL), scrambled negative control (NEG), or toxicity positive control (TOX) ASOs were delivered gymnotically to MPCs at up to three doses (0.25 µM, 0.5 µM, 1.0 µM) on D0, and myotubes were analysed at D5 of differentiation. **A** MAL transcript knockdown was quantified by qPCR. **B/C**. Cell viability was analysed with Cell Titre Glo 2.0 ATP assay. **D.** Day 5 myotubes were fixed and stained for F-actin (phalloidin, red), pan-myosin (MF-20, green) and nuclei (DAPI, blue). A representative image (four central images from a single well stitched together) is shown for each line. Scale bars are 200 µm. **E/F**. As a measure of myotube morphology following ASO treatment, the percent myosin area in the whole well was calculated. Data are mean±SEM of *n* = 3 independent replicates, and data were normalised to the NEG treated myotubes. Statistical comparisons are unpaired t-test with significance set at *p* < 0.05. Welch’s correction was performed when the variance between groups was significantly different.

Force-frequency relationships of p.R2508C tissues were also affected: unlike control tissues (which showed a mostly sigmoidal force-frequency curve), p.R2508C tissues required higher stimulation frequencies to reach maximal tetanic force, resulting in a more linear curve (Fig. 4M). Although this was observed at both D14 and D21, it was more pronounced at day 14 where both controls achieved >50% maximal force output at 1 Hz, whereas p.R2508C required 5 Hz to reach 50% of maximal force (Fig. 4M). At D14, maximum force (as a percentage) of p.R2508C tissues was significantly reduced compared to controls at all stimulation frequencies up to and including 20Hz (Fig. 4M, *p* < 0.05), while no significant differences were observed between the two controls (Fig. 4M, *p* > 0.05). Altogether, these data show a clearly altered force-frequency response and imply that deficits of p.R2508C tissues may be more pronounced at lower stimulation frequencies.

### RYR1 patient derived myotubes are a useful model for screening RNA-modifying therapeutics

Having established our cell lines as a useful *RYR1*-RM disease model, we next sought to determine whether *RYR1*-RM patient derived myotubes might be a tractable testbed for screening gene-targeting therapeutics. To this end, we aimed to establish methods for medium-and high-throughput delivery of therapeutics and associated downstream analyses of efficacy and toxicity. As an exemplar, we chose to explore delivery of antisense oligonucleotides (ASOs) since these can be used to achieve potent and selective knockdown of pathogenic mRNA^73^, a strategy previously shown to be feasible in two mouse models harbouring dominant pathogenic variants in *RYR1*^35^. As a positive control, we selected a 5’FAM-labelled and LNA-modified ASO targeted to the long noncoding RNA *MALAT1* (MAL). As a negative control, a scrambled ASO with no known targets in the human transcriptome was used (NEG).

Initially, ASOs at high concentration (1.0 µM) were delivered gymnotically to healthy control (CTRL-3) and patient-derived (p.R2508C) MPCs with differentiation media at D0. Myotubes were subsequently analysed for target transcript (*MALAT1*) knockdown at D5 of differentiation. At this dose, ASO uptake was highly efficient, with FAM detected in almost 100% of p.R2508C nuclei (99.91% ±0.10%). This highly efficient ASO uptake resulted in more than 90% *MALAT1* transcript knockdown compared to NEG-treated samples in both CTRL-3 and p.R2508C myotubes (Fig. 5A; CTRL-3, *n* = 5, *p* < 0.0001; p.R2508C, *n* = 3, p = 0.0016, students unpaired t test with Welch’s correction).

To establish assays for cell viability we used a positive control ASO known to induce cellular toxicity (TOX) at a wider range of doses (0.25 µM, 0.5 µM, 1.0 µM). We assessed cell viability using an ATP assay (CellTiter-Glo® 2.0), and by quantifying myosin area. Using these workflows, we could detect reductions in cell viability as little as 10% in the 1.0 µM treatment condition (Fig. 5B, CTRL-3, *n* = 3, *p* = 0.01; Fig. 5C, p.R2508C, *n* = 3, *p* = 0.04, unpaired t test) relative to the NEG. Changes to myotube morphology (Fig. 5D) and a significant reduction in total myosin area (Fig. 5E; CTRL-3, *n* = 3, *p* = 0.046; Fig. 5F, p.R2508C, *n* = 3, *p* = 0.016, unpaired t test) in CTRL-3 and p.R2508C myotubes were also observed relative to NEG treated cells. Together, these results suggest that the established workflows are useful for identifying lead candidate ASO therapeutics with suitable on-target activity and low cellular toxicity, confirming the utility of these iPSC-derived skeletal muscles as a therapeutic screening platform in 2D culture.

## Discussion

This study sought to establish the utility of patient-iPSC-derived myogenic progenitor cells for disease modelling and screening of targeted therapeutics in the context of dominant *RYR1*-RM. These patient cell resources fill a gap in available *RYR1*-RM disease models by creating a potentially unlimited source of patient material, that retains the patient’s genomic context and enables cell and tissue-specific investigations. Importantly, we have shown that when differentiated to 3D engineered muscle tissues, these cells can recapitulate key features of disease, including force deficit and abnormal relaxation kinetics. These data justify future investigation of additional patient lines as well as expansion of the possible variants, assays and phenotypes that could be studied.

Here, we created patient-derived MPCs from five unrelated patients harbouring dominant pathogenic variants in the *RYR1* gene who had a diagnosis of CCD with (p.V2168M, p.R2508C) or without (p.N1346K, p.4715-18del, p.H4813Y) concurrent MHS. Patient-derived cells were compared to multiple independent healthy control lines. A limitation of cell-based studies is that underlying differences in the cell populations (e.g. variable proportions of different cell types) can confound interpretation of results^74^. Thus, we sought to comprehensively profile the myogenic progenitor populations we created before proceeding to phenotypic analyses. Our qPCR panel data support the ability of the directed differentiation procedure to create a mixture of biologically relevant cell types including early myogenic progenitors (*SIX1, PAX3*), mesenchymal progenitors (*PDGFRA, PRRX1)* and fibroblasts (*DCN*). Fusion index data and staining for RyR1 support the conclusion that our five patient lines are similar in composition and maturity and are within the range of the two best performing healthy control lines (CTRL-2, CTRL-3) for all parameters. One healthy control line (CTRL-1) was an outlier which had much lower proliferation rate and fusion index. We opted to include these data as an illustration that poor fusion cannot be assumed to be associated with a patient-phenotype, and as a cautionary note to ensure that adequate consideration is given to selection of appropriate healthy control lines for each patient line or batch of differentiation experiments.

Having establish the capacity of patient-derived MPCs to produce high quality myotubes in 2D, we next investigated their utility for disease modelling. Two of the substitutions (p.V2168M, p.R2508C) are recurrent and have previously demonstrated increased sensitivity to caffeine accompanied by increased resting cytosolic [Ca^2+^] and reduced maximal [Ca^2+^] release following stimulation with caffeine in HEK cells *in vitro*^28,29^. The p.R2508C variant was also associated with accelerated Ca^2+^-induced Ca^2+^ release (CICR) rate^15,54^. These *in vitro* studies shed light on the potential mechanisms but lack human muscle-cell context. Here, we focussed our proof-of-principle study on the p.R2508C substitution, which has been extensively investigated, and was associated with the earliest onset of disease symptoms in our cohort. In line with a previous study^28,29^, using a 2D whole-well Ca^2+^ assay, we were able to show that relative to healthy control myotubes, p.R2508C myotubes produced significantly reduced total Ca^2+^ release following stimulation with caffeine. This important finding confirms that these cells can produce Ca^2+^ transients in response to caffeine and suggests that this medium-high throughput assay produces biologically meaningful outputs.

We do acknowledge that caution should still be taken in the interpretation of whole-well data. We have used the most closely matched healthy control and patient cell lines for this assay in terms of myogenic potential (fusion index and RyR1 abundance), but despite this, we cannot rule-out the possibility that there may be underlying differences in the cultures (such as maturity) that contribute to the difference in total Ca^2+^ release. In future, normalisation to fusion index, myotube area or RyR1 abundance could be performed to further validate comparisons between different cell lines or experiments. In our hands, the necessity of culturing the myotubes on a hydrogel for the Ca^2+^ assay to mitigate detachment, prevented precise quantification of these metrics, as hydrogel thickness precluded imaging using our current workflows. However, even without normalisation, this assay will be useful for therapeutic screening where maximal Ca^2+^ release is compared within a single line before and after drug treatment and therefore myogenic potential is internally controlled. The benefit of the whole-well application in this context is the increased ease and scalability, although this may come at the cost of increased noise (compared to traditional single myofibre measurements). Additional parameters such as dose-response could also be analysed in future.

Ultimately, we aimed to use these cell lines to model patient-relevant phenotypes such as muscle weakness. To achieve this, we employed the Curi Bio Mantarray^TM^ platform for generating 3D EMTs which enabled non-invasive, longitudinal monitoring of contractile performance in real-time^75^. This system has previously been used to model functional deficits in force output (twitch force, tetanic force, contraction velocity etc.) in other disease contexts^75^. These studies also demonstrated that healthy control EMTs produce increased contractile force when exposed to low doses of ryanodine (known to induce the RyR1 open state), while contractile responses were completely abolished in the presence of high doses of ryanodine (known to force the RyR1 closed state)^75^. These experiments demonstrated that RyR1 functional state is directly linked with contractile performance in the Mantarray^TM^ and established this platform as suitable for modelling RyR1 dysfunction.

Using the p.R2508C variant line as an exemplar, we show a striking and significant reduction in both twitch and tetanic force output, accompanied by increased time to peak and delayed relaxation in patient-derived 3D EMTs. Qualitatively, we also observed an increase in passive tension of EMTs over time, which is concordant with findings from a homozygous p.R2509C mouse model in which sarcomere shortening was reported^76^. Together, the increase in passive tension and impaired relaxation imply that the p.R2508C variant leads to Ca^2+^ leak, likely via destabilisation of RyR1 structure which favours the open state. This is supported by data generated in HEK cells which show the p.R2508C variant exhibited both decreased peak [Ca^2+^] release and elevated resting [Ca^2+^]^29^. Indeed, in our hands, p.R2508C myotubes grown on a 2D hydrogel at days 14-20 exhibited significantly reduced Ca^2+^ release, which aligns with the reduced force output and disrupted relaxation kinetics observed in the 3D EMTs during the same timeframe. Thus, we have been able to directly link perturbed RyR1 channel function to downstream disruption of muscle contractile performance.

A limitation of our 3D tissue studies is the small number of tissues we have analysed. For example, a single EMT per cell line per time-point was used for analysis of effective CSA. Ideally, each individual trace would be normalised to CSA of the matched EMT, however, this severely restricts possible downstream analyses (such as proteomic studies, which are ongoing). As a proof-of-principle, we performed normalisation of force traces to matched CSA for each of our three cell lines at two time points, which demonstrated that while the absolute values varied, the overall interpretation of the data was unchanged by the normalisation. While this does require further validation, it raises the possibility that normalisation may not be necessary if the initial starting cell populations are similar between groups (e.g. as determined by cell composition and fusion index). As yet, there are no field-wide gold standards specified for EMT analyses such as standardisation of pacing routines, analyses time-points, CSA analyses, and use of mixed or FAC-sorted cell populations^77^. This presents a challenge as force values and phenotypic outputs cannot be easily compared between studies.

Future studies will model the remaining four patient lines in 3D as well as an expanded set of potential phenotypes and in-depth profiling by proteomics and metabolomics. Furthermore, we will establish *RYR1*-RM 3D EMTs as a useful treatment screening platform. In the context of neuromuscular disorders, 3D culture models have already been used to screen the effectiveness of small molecule drugs^78^ and rAAV9-mediated gene therapy^79^, although neither study was able to show rescue of muscle force-deficits following treatment. Since contractile force is a key outcome measure of many muscle diseases, the ability to quantify improvements in contractile performance would be a powerful predictor of treatment efficacy. Additional development and testing of treatments in 3D EMTs will be needed to realise their full potential as a pre-clinical screening tool.

A crucial outcome of our study was optimisation of methods to deliver LNA-modified ASOs and screen their therapeutic knockdown effects and toxicity using high-throughput assays in 2D myotube cultures. We show that gymnotic delivery of LNA-modified ASOs results in highly efficient uptake into the nucleus, commensurate with strong knockdown and minimal cellular toxicity. These workflows could be readily adapted to screening other therapeutic modalities in *RYR1*-RM myotubes or in alternative disease contexts. Ultimately, as part of treatment development workflows, we propose that lead candidate therapeutics identified in medium-high throughput 2D culture, could be prioritised for further screening of functional benefit in the 3D EMT model.

In sum, we report the differentiation and characterisation of myogenic progenitor cells from five unrelated *RYR1*-RM patients with pathogenic variants spread throughout the length of the RyR1 protein. We establish the quality of these cell resources and describe methods for the using these cell lines as a treatment screening platform in 2D myotubes. Further, we present evidence that the Mantarray^TM^ 3D EMT platform is a useful tool for modelling of *RYR1* patient-relevant phenotypes. To the best of our knowledge, this is the first report to show a functional deficit in muscle contractile performance in 3D EMTs generated from an *RYR1*-RM patient. Together, the patient-derived cell lines we have developed represent a useful resource for the *RYR1*-RM community.

## Methods

### Human samples, consent and ethics

The use of human cells in this study was approved by the University of Western Australia’s Human Research Ethics Committee (approval number: RA/4/20/1008) and the Child and Adolescent Health Service Human Research Ethics Committee (approval number RGS0000004546, ‘The Perth Neurogenetic Biobank for Genetic and Functional Genomics Research’ (supersedes 2019/RA/4/20/1008)), which was recognised by the UWA human ethics office. Some cell lines were obtained with consent from the National Muscle Disease Bio-databank (NMDB), approved by The Royal Children’s Hospital Melbourne Human Research Ethics Committee (HREC Project Number: 90530), or from the Genethon cell bank (activity authorization No. AC-2018-3156, import/export authorization No. IE-2018- 994) with signed consent for genetic testing, banking and research.

Skeletal muscle biopsies (*vastus lateralis*) were obtained as excess from *in vitro* contracture testing with informed consent for research purposes (Murdoch University Human Ethics Committee, 2017/101).

### Protein structure

The crystal structure of rabbit RyR1 in complex with FKBP12 at 3.8 Å (PDB ID: 3J8H^80^) was interrogated using the RCBS Protein Data Bank (RCSB.org, accessed 2^nd^ September 2025)^81^ and was visualised and colourised using Mol*^82^. Human RyR1 amino acid numbering has been applied to the rabbit structure according to the protein sequence <u>NP_000531.2</u>.

### Culture and directed differentiation of induced pluripotent stem cells (iPSCs)

All iPSC lines used in this study were previously verified for genome stability and trilineage differentiation capacity and were cultured as previously described^49–51^. Healthy control iPSCs were obtained from various sources: CTRL-1 (iPSC-HC01-5) was a gift from Prof. Eric Olson, CTRL-2 (iPSC-GM23720) was purchased from the Coriell cell repository, and CTRL-3 (iPSC-SCTi003-A) was purchased from StemCell Tech. iPSCs were differentiated to muscle progenitor cells (MPC) using the STEMdiff™ Myogenic Progenitor Supplement Kit (Stem Cell Technologies) for 30 days according to the manufacturer’s instructions. Additional details are provided in the Supplementary Information.

### Culture and terminal differentiation of myogenic progenitor cells (MPCs)

MPCs were cultured in complete MyoTonic^TM^ medium (MyoTonic^TM^ basal medium (Cook Myosite) supplemented with MyoTonic^TM^ serum-free growth supplement (Cook Myosite), 10% FBS (Gibco, Australian Origin), 1% Pen/Strep (in-house) and 20 ng/mL FGF2 (PeproTech, Cat#100-18B)) with media changed every 2-3 days and maintained below 80% confluence. At 60-80% confluence, MPCs were harvested with TrypLE^TM^ Express and subcultured at a density of 0.5-1 × 10^4^ cells/cm^2^. For secondary differentiation to myotubes, MPCs were plated at 2 × 10^4^ cells/cm^2^ on Matrigel (standard formulation (Cat# 354234), 1:100 in DMEM/F-12; Thermo Fisher) in complete MyoTonic^TM^ until they reached 85-95% confluence. Once suitably confluent, media was replaced with myotube differentiation media (DMEM/F-12 (in-house), 1% N2 supplement (Gibco, Cat#17502-001), 1% ITS-X (Gibco, Cat#51500-056), 1% GlutaMAX^TM^ (Gibco), 1% P/S (in-house)) for up to 7 days, changing media every four days.

### RNA extraction and cDNA synthesis

RNA was extracted using the RNeasy Plus Mini (for ∼1 × 10^6^ cells in suspension) or Micro (for scraping from a 12-well plate) kits (QIAGEN) according to the manufacturer’s instructions. Homogenisation was performed using a QIAshredder column (QIAGEN) and RNA was eluted using UltraPure dH2O (Gibco). For extraction of RNA from primary human tissue, QIAzol reagent (QIAGEN) was used according to the manufacturer’s protocol with minor modifications to the homogenisation step. Briefly, vastus lateralis muscle tissue (∼30-50mg) was homogenised in 1 mL QIAzol reagent in a tough-tube containing 2-3 ceramic beads using the Mini-Beadbeater-24 (BioSpec Products) at 2.75 oscillations/second for 30 seconds. RNA was eluted in 30-50 µL of UltraPure dH2O. RNA concentration, purity, and quality were assessed using a NanoDrop One Spectrophotometer (ThermoFisher) and the 4200 TapeStation system using RNA ScreenTape and reagents (Agilent Technologies). cDNA was synthesised using the LunaScript® RT SuperMix Kit (NEB) according to the manufacturer’s instruction.

### RT-qPCR

RT-qPCR was performed using the QuantiNova SYBR Green PCR Kit (QIAGEN) on a QuantStudio 6 Pro Real-Time PCR System (Applied Biosystems). The following cycling conditions were used: 95 °C/2 min, (95 °C/5 s, 60 °C/10 s) × 40 cycles, followed by melt curve analysis. Data were analysed using the Thermo Fisher Connect Platform. CT values were obtained by applying a constant ΔRn threshold of 0.2 and analysis performed using the 2^-ΔCT method, normalised to the geometric mean of the CT values for *TBP* and *EEF2* (endogenous control genes). Primers used are listed in Supplementary Table S4. Some primer sequences were sourced from the others^60,83^.

### Immunocytochemistry

Media was removed and cells were washed twice with 1X DPBS prior to fixing in 4% PFA (in DPBS) for 15 min, followed by washing three times with 1X DPBS. Cells were permeabilised with 0.2% Triton X-100 in DPBS for 15 min at RT, washed twice with 0.2% Triton X-100 then incubated in blocking buffer (3% BSA + 0.2% Triton X-100 in PBS) for 60 min at RT with rocking. The appropriate stain, primary and/or preconjugated antibodies (Table 2) were diluted in blocking buffer and incubated at 4 °C overnight with rocking. Secondary antibodies (Table 2) were diluted in blocking buffer and incubated for 1 h at room temperature on a rocker. NucBlue Fixed Cell Ready Probes Reagent (Invitrogen) was diluted 2 drops per mL in DPBS and incubated for 10 min, after which the cells were washed once more with PBS. Plates were imaged using a CellInsight CX7 High Content Analysis Platform (Thermo Scientific).

**Table 2.**
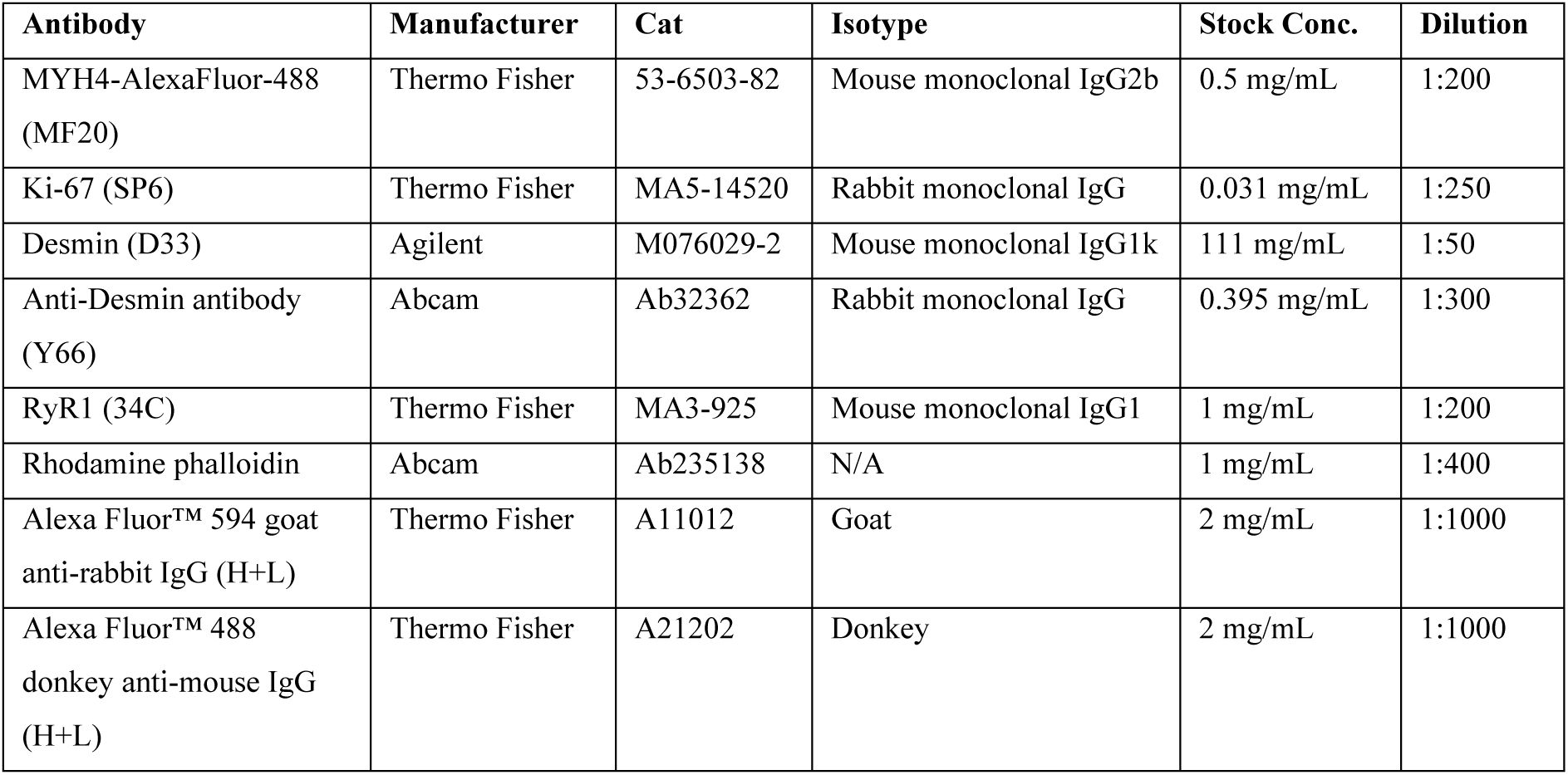
Antibodies and stains used in this study.

| Antibody | Manufacturer | Cat | Isotype | Stock Conc. | Dilution |
| --- | --- | --- | --- | --- | --- |
| MYH4-AlexaFluor-488 (MF20) | Thermo Fisher | 53-6503-82 | Mouse monoclonal IgG2b | 0.5 mg/mL | 1:200 |
| Ki-67 (SP6) | Thermo Fisher | MA5-14520 | Rabbit monoclonal IgG | 0.031 mg/mL | 1:250 |
| Desmin (D33) | Agilent | M076029-2 | Mouse monoclonal IgG1k | 111 mg/mL | 1:50 |
| Anti-Desmin antibody (Y66) | Abcam | Ab32362 | Rabbit monoclonal IgG | 0.395 mg/mL | 1:300 |
| RyR1 (34C) | Thermo Fisher | MA3-925 | Mouse monoclonal IgG1 | 1 mg/mL | 1:200 |
| Rhodamine phalloidin | Abcam | Ab235138 | N/A | 1 mg/mL | 1:400 |
| Alexa Fluor™ 594 goat anti-rabbit IgG (H+L) | Thermo Fisher | A11012 | Goat | 2 mg/mL | 1:1000 |
| Alexa Fluor™ 488 donkey anti-mouse IgG (H+L) | Thermo Fisher | A21202 | Donkey | 2 mg/mL | 1:1000 |

### Image analysis and quantification using CellProfiler

Image analysis was performed using custom CellProfiler pipelines. Nuclei were detected using an Otsu threshold with segmentation based on intensity. To calculate fusion index, a binary mask representing the myotubes was generated using a manual threshold of MF20 signal, then applying steps to fill areas of low signal where nuclei were located. Nuclei were classified as within myotubes if ≥ 50% of their area overlapped with the binary myotube mask. Fusion index was expressed as the total area of the nuclei within myotubes as a percentage of the total area of all nuclei in each well. Ki67-positive nuclei were classified using a manual threshold on the Ki67 channel within the area of each nucleus. Nuclei within desmin-positive myotubes were classified using a manual threshold of the desmin channel around the edge of each nucleus. To quantify RyR1 expression, a binary myotube mask was generated as described above, and the total RyR1 signal within the boundaries of the mask was calculated for each well.

### Gymnotic delivery of ASOs to myotubes

MPCs were cultured and plated for terminal differentiation as described above. On D0 growth media was replaced with freshly prepared myotube differentiation media supplemented with ASOs to a final concentration of 0.25 µM, 0.5 µM or 1.0 µM. ASOs were added to each plate in technical duplicate. Media was not changed, and plates were harvested at D5 of differentiation.

To analyse transcript knockdown, cell lysates and cDNA were generated using the PowerSYBR^TM^ Green Cells-to-CT^TM^ kit (Invitrogen, Cat#4402955) according to the manufacturer’s instructions. To assess cell viability, we used the CellTiter-Glo® 2.0 Cell Viability Assay (Promega) per manufacturer’s instructions. Luminescence readings were recorded using the CLARIOstar plate reader (BMG Labtech). Myosin area was determined by imaging on the CellInsight CX7 High Content Analysis Platform (Thermo Scientific) as described above. Additional details, include full ASO sequences, are provided in Supplementary Information.

### Hydrogel fabrication

A solution of 5% (w/v) gelatin (from porcine skin, gel strength ∼175 g Bloom, Type A; Merck, Cat# G2625) was made using 65 °C PBS (Gibco). A 20 U/mL solution of transglutaminase (Moo Gloo TI, Modernist Pantry) was made using 37 °C PBS. Both solutions were sterilised using a 0.45 µm filter. These two solutions were mixed 1:1 and cast into 96-well plates at 32 µL per well (100 mL/cm^2^). Plates were incubated at 4 °C for 15 min to set and then at 37 °C in an incubator for 4 h to cross-link. Hydrogels were then washed twice with PBS and once with MyoTonic^TM^ Complete media (10 min per wash). Hydrogels were stored in MyoTonic Complete media overnight at 37 °C in an incubator before seeding cells.

### Whole well calcium assay

MPCs were seeded onto hydrogel-coated 96-well plates and differentiated as described above. N2 differentiation media was replaced every 96 hours. Ca^2+^ flux was measured using the Fluo-4 Direct Calcium Assay Kit (Invitrogen, F10471). The Fluo-4 probe was suspended in 10 mL of Hanks’ Balanced Salt Solution (HBSS). Prior to the assay, N2 media was removed, and cells were washed once with HBSS (100 µL/well). HBSS was replaced with 50 µL fresh HBSS and 50 µL Fluo-4 solution. Cells were protected from light and incubated at 37 °C for 1 h. The Fluo-4 solution was removed and replaced with 20 µL HBSS per well. The plate was inserted into a CLARIOstar plate reader (BMG Labtech) at room temperature. Fluorescence data was captured using a spiral imaging pattern within a 2 mm area in the centre of the well, from a focal distance of 4 mm. Detector gain was set to 2500. Data was captured every 0.5 s for 30 s, then 20 µL of caffeine solution was injected at 100 µL/s before resuming data capture for a further 30 s.

### Generation of 3D engineered muscle tissues (EMTs)

#### Preparation

3D engineered muscle tissues (EMTs) were prepared using the Curi Bio Mantarray^TM^ platform. Casting posts were coated with 0.1% PEI (Sigma, #P3143) for 10 min at RT, rinsed in sterile dH2O for 5 min, then coated with 0.01% glutaraldehyde (Sigma, 111-30-8) for 30 min at RT, washed twice for 5 min with sterile dH2O and left to dry. Casting plates were pre-prepared by chilling at 4 °C for at least 30 min, then pre-coated with diluted thrombin (3 µL of 100 U/mL thrombin + 47 µL DMEM per well), ensuring even coverage within each casting trough. Thrombin-coated casting plates were used within 3 h of preparation. Prior to casting, MPCs were cultured in “skeletal muscle growth medium” (skGM): DMEM high glucose with pyruvate (Gibco, #11995-065), 10% FBS (Hyclone, #SH30071.03HI), 1 µM dexamethasone (Sigma, #D4902), 100 nM insulin (Lonza, #BE02-033E20), 40 ng/mL FGF2 (R&D, #3718-FB-100). Normal human dermal fibroblasts (NHDF) were sourced from Lonza (Cat# CC-2511) and cultured in DMEM high glucose, pyruvate (Gibco, #11995-065) with 10% FBS (Hyclone).

#### Tissue casting

MPCs (∼P4, ∼70-90%) and NHDF fibroblasts (∼P5, ∼80-90%) were harvested using 1X TrypLE, resuspended in DMEM and counted using a Countess II automated cell counter (Thermo Fisher). MPCs were resuspended to 750,000 cells per 30 µL and fibroblasts to 75,000 cells per 30 µL. A mastermix was prepared for 12x EMTs plus 20% excess, to result in the following quantities per individual EMT: 30 µL MPCs, 30 µL fibroblasts, 30 µL GFR Matrigel (neat, Cat# 356231, Lot# 2318004), and 10 µL fibrinogen (50 mg/mL stock). 100 µL of cell mixture was carefully added to each casting well (pre-coated with thrombin, chilled on ice), then triturated five times per well. All casting steps were performed on ice. Casting plates were then transferred to a 37 °C incubator for 80 min, after which time 1 mL of MyoTonic growth medium + 2 g/L amino caproic acid (ACA, Thermo Fisher, #103301000) was added to each well and incubated at 37 °C overnight (20 h). Then, the post lattice (with EMTs attached) was carefully transferred to a pre-prepared 24-well plate with 2 mL/well of pre-equilibrated EMT differentiation medium (DMEM high glucose with 1X NEAA (Gibco, #11140050), 2% horse serum (Gibco #16050-114), 1X B27 supplement (Gibco #17504-044), 10 ng/mL IGF1, 10 ng/mL HGF, 10 µM SB431542, 2 g/L ACA). Medium was changed every other day. From day 10 onwards, ACA concentration was increased to 5 g/L to mitigate excessive tissue remodelling.

### EMT stimulation, data recording and analysis

Electrical stimulation of EMTs was commenced on day 6 and continued until day 21 (endpoint). Recordings were taken for each stimulation. In brief, each day EMTs were sequentially measured as follows: a) 2 min with no stimulation to assess baseline and detect any appreciable spontaneous contractions, b) 2 min of 1 Hz (twitch) protocol, and c) force-frequency protocol (1 - 50Hz). Detailed information about stimulation protocols can be found in Supplementary Methods. Stimulation data were all collected using the Mantarray controller software and analyzed using the Pulse 3D cloud analysis platform (v1.0.8), using analysis methods recommended by Curi Bio.

### EMT harvesting and immunostaining

On harvest days (D7, D14 and D21), tissues were rinsed in cold 1X DPBS and either a) flash frozen in liquid nitrogen (for RNA/protein work), or b) rinsed in cold 1X DPBS, then fixed in 4% PFA overnight at 4 °C (for immunostaining). Following immunostaining and confocal imaging, cross-section images were analysed using a custom ImageJ script with manual outlining of tissues based on staining borders using the ‘freehand’ tool, followed by calculation of cross-sectional area and myosin area (MF20-positive pixels as a percentage of total pixels). These values were used for normalisation of twitch force measures for matched tissues. Detailed methods are available in Supplementary Information.

### Figures, graphs and statistics

Graphs and statistics were generated using GraphPad Prism version 10.5.0 for Windows, (GraphPad Software, Boston, Massachusetts USA, www.graphpad.com). Schematic figures were created using BioRender.

## Acknowledgements

This work was supported by research grants from the Channel 7 Telethon Trust to CI Taylor (2024, 2025) and by a research grant from The Stan Perron Charitable Foundation (APP-202003010Research, PI Taylor). This work was also supported by The Children’s Health and Disability Foundation WA and Hearts & Minds Investments.

JSC is supported by a Harry Perkins Institute of Medical Research Safe Harbour Fellowship. This research was supported by the Commonwealth through an Australian Government Research Training Program Scholarship [DOI: https://doi.org/10.82133/C42F-K220] to JJC. RLT is supported by a Harry Perkins Institute of Medical Research Safe Harbour Fellowship.

VC and PH are supported by the MRFF 2021 EPCDR Initiative (Chronic Musculoskeletal conditions in children and adults – Application ID 2015993).

GR is supported by an Australian NHMRC Investigator Grant (APP2007769) and the Patricia Verne Kailis Fellowship.

We would like to acknowledge the advice and guidance provided by E/Prof Sue Fletcher, who also provided access to human muscle biopsy samples. We would also like to acknowledge the thoughtful discussion of *RYR1* variants, and their analysis, provided by Prof Susan Treves.

