## Supplementary Figures and Methods for "Ryanodine receptor 1 (*RYR1*) patient-derived muscle cells recapitulate disease phenotypes in 2D and 3D culture models"

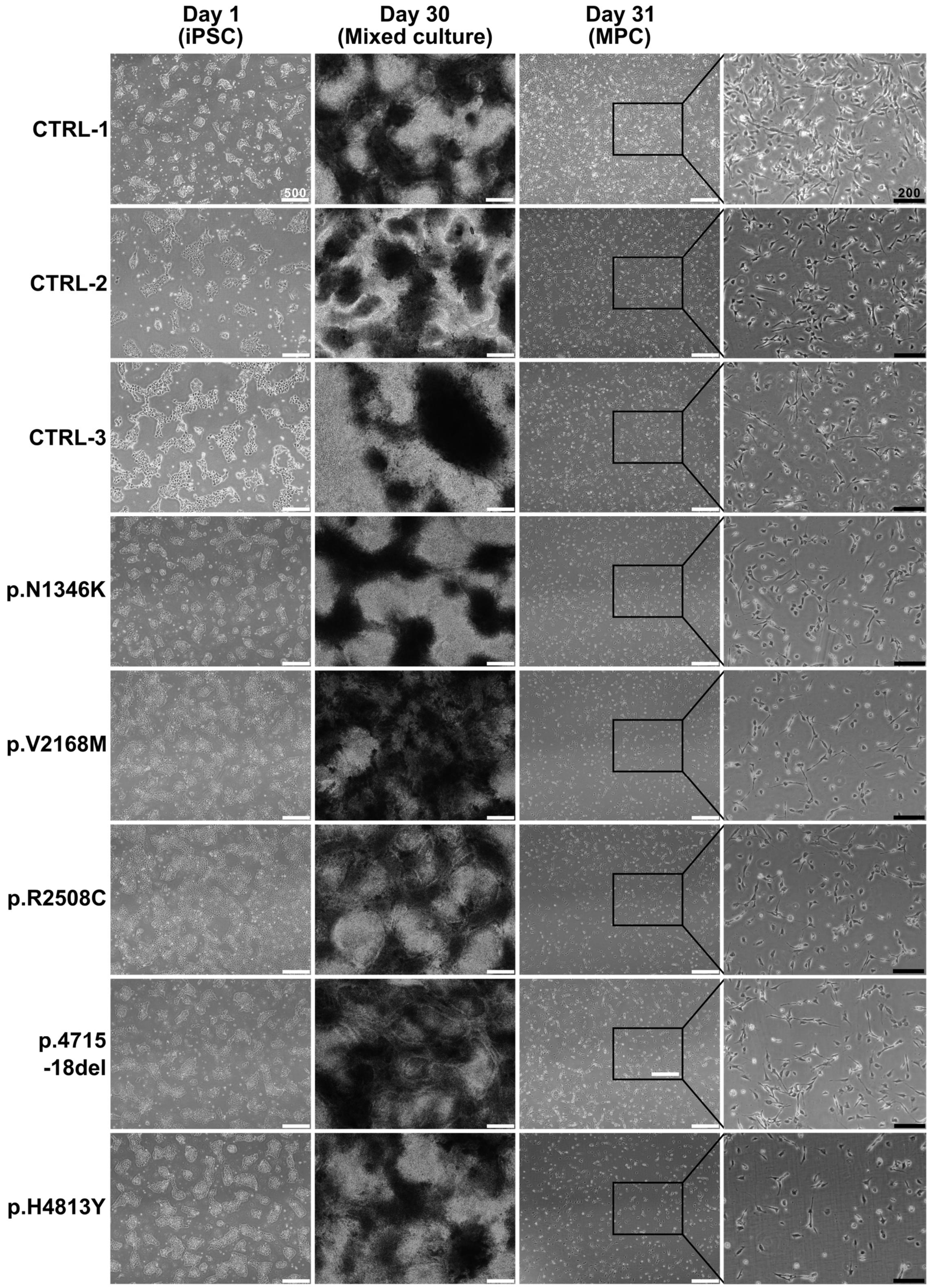


**Supplementary Figure S1. Brightfield microscopy of cells during the iPSC to skeletal muscle directed differentiation.**

Representative brightfield images of cells at day 1 of differentiation (iPSCs), day 30 of differentiation (dense, mixed culture) and following dissociation to single cells (MPC, myogenic progenitors). Representative images from *n* = 3 independent differentiations are shown, except MPC-4715-18del which is *n* = 2. White scale bars are 500 µM, black scale bars are 200 µm.

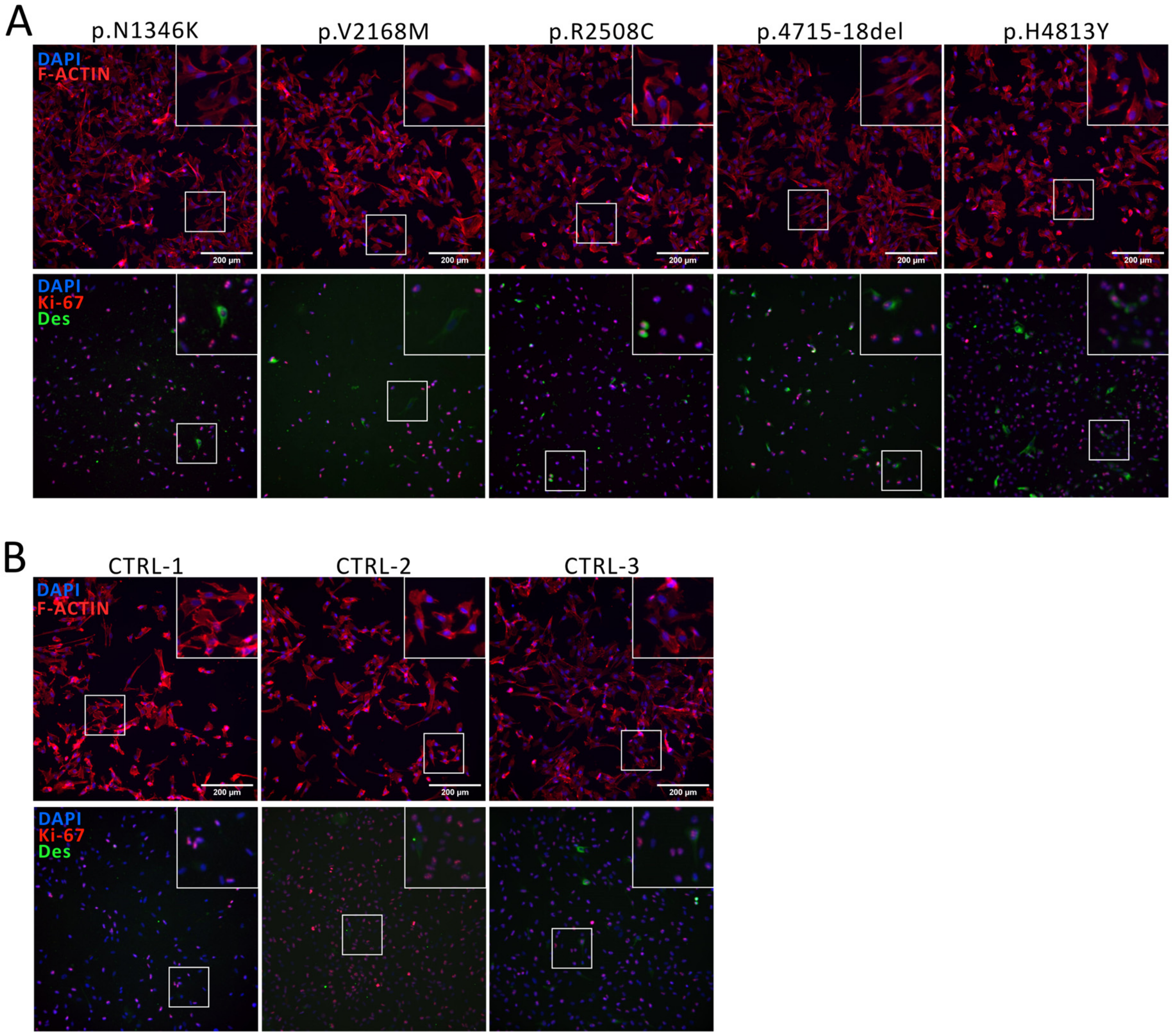


**Supplementary Figure S2. Staining of myogenic progenitors to assess morphology, proliferative status and myogenic lineage state.**

Myogenic progenitors (MPCs) from (**A**) *RYR1*-RM patients and (**B**) healthy controls were fixed and stained for (top panels) filamentous actin (F-Actin, red, cytoplasmic localisation), or (bottom panels) Ki67 (red, nuclear localisation) and desmin (green, cytoplasmic localisation). All wells were counterstained with DAPI (blue) to identify nuclei. Representative images from $n$ = 3 independent differentiations are shown, except MPC-4715-18del which is $n$ = 2. Scale bars is 200 µm and applies to all images

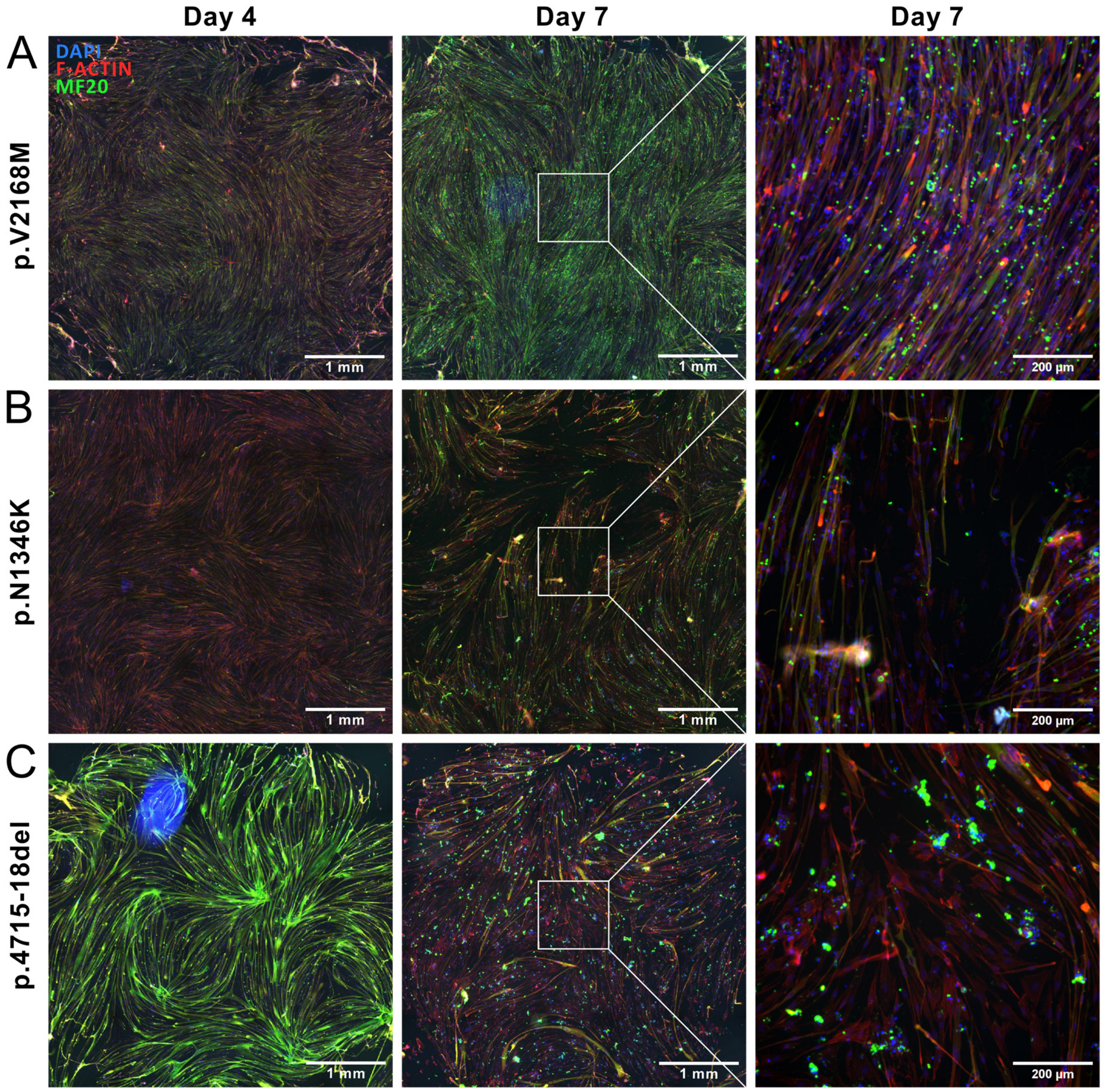


**Supplementary Figure S3. Variable myotube loss by Day 7 of myotube differentiation.**

**A.** Example of a line with no peeling at Day (D) 7. **B.** Example of a line with localised patches of peeling at D7. **C.** Example of a line with majority of the well peeling at D7. Left and centre images are full 96-well images composed by stitching 25-fields of view together. Scale bars are 1 mm. Inset (right) is a single 10x magnification image from the centre of the 96-well. Inset scale bars are 200 µm.

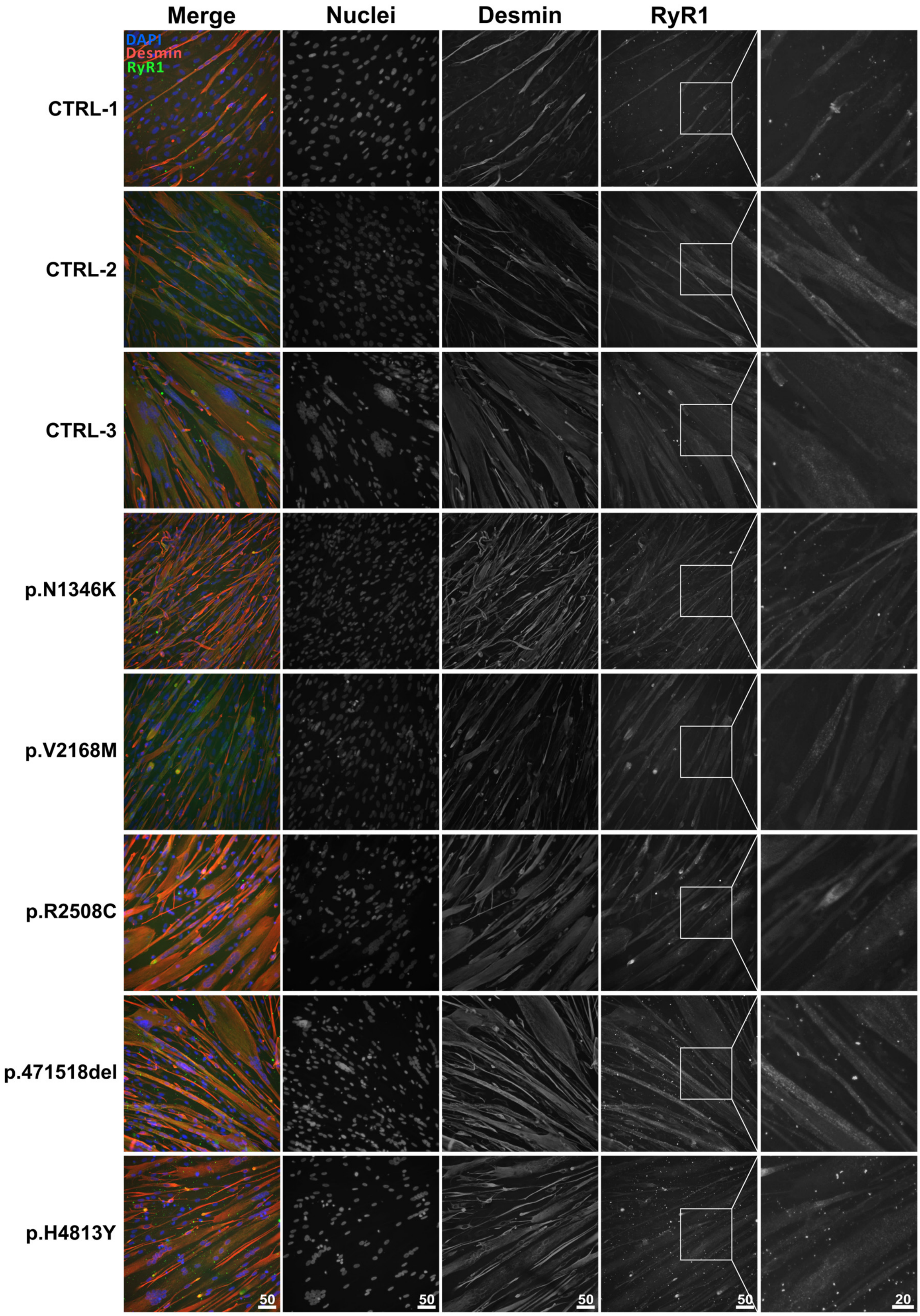


**Supplementary Figure S4. Myotubes stain positively for RyR1 at Day 4 of differentiation.**

Myotubes fixed at Day 4 were stained with DAPI (nuclei, blue and second column), desmin (red and third column) and RyR1 (34C, green and fourth column). Scale bars are 50 µm, except for inset (right column) which are 20 µm.

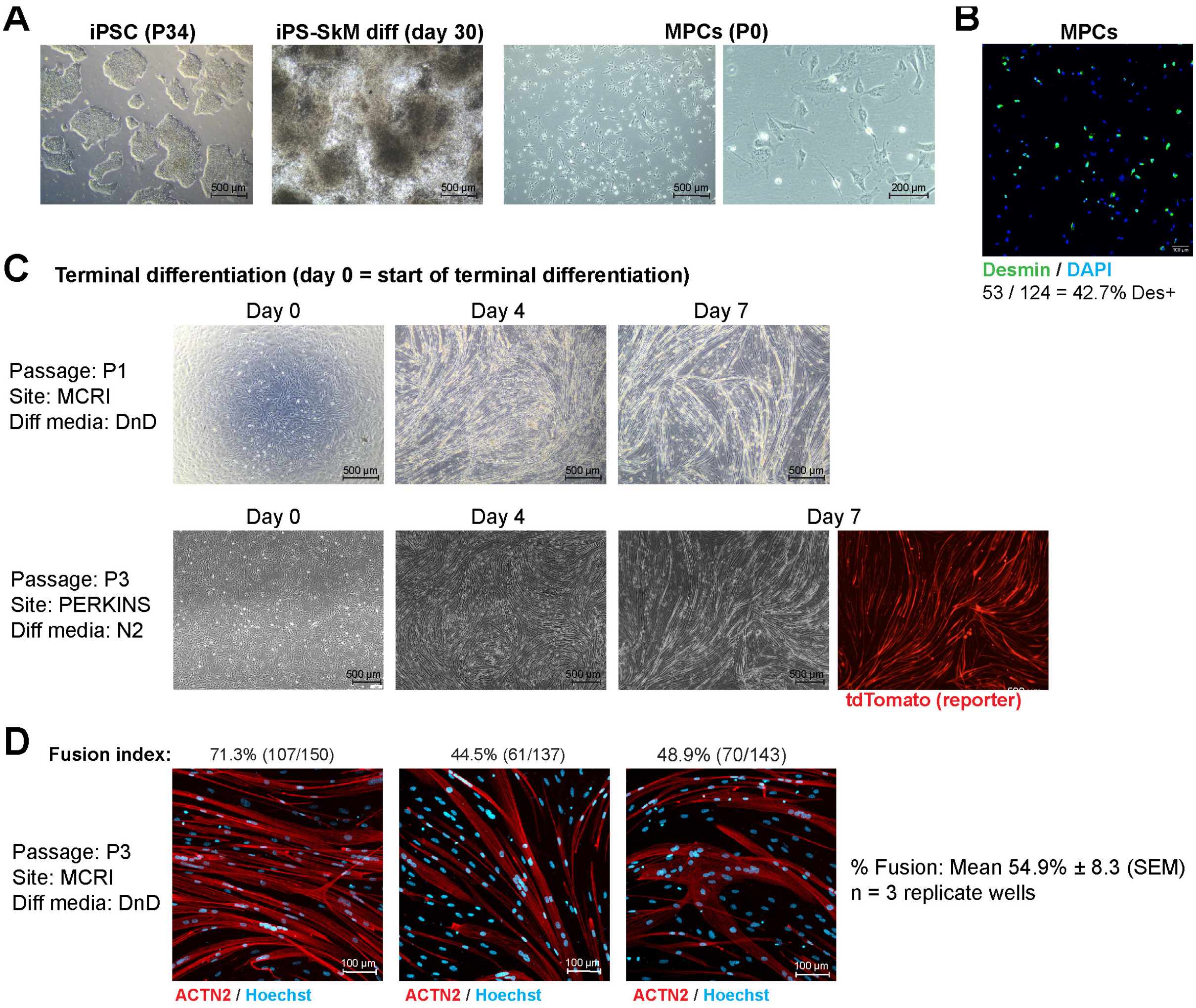


**Supplementary Figure S5. Characterisation of CTRL-4 (ACTA1-tdTomato) control MPCs.**

**A)** Brightfield images of iPSCs (passage 34), iPSCs at day 30 of STEMdiff™ skeletal muscle differentiation (iPS-SkM diff) and resulting myogenic progenitor cells (MPCs) after differentiation harvest (passage 0). **B)** Desmin staining of MPCs. **C)** Terminal differentiation to assess fusion of MPCs in 2D. Top: differentiation of P1 MPCs at MCRI site (using in-house DnD differentiation media; α-MEM media with 1% pen/strep, 0.5% ITS, 2% B-27 supplement, 10 µM DAPT, 1 µM dabrafenib). Bottom: differentiation of P3 MPCs at primary study site (PERKINS), differentiated using N2 media (same as used for other lines in the paper; DMEM/F12 GlutaMax with 1% N2 supplement, 1X ITS-X). **D)** Immunostaining of myotubes at day 4 of terminal differentiation stained for ACTN2 (red; myotubes, #ab68167) and Hoechst (blue; nuclei). Fusion index is indicated above each image (mean 54.9 ± 8.3%, *n* = 3 images from replicate wells).

**Supplementary Methods**

*Culture and directed differentiation of induced pluripotent stem cells (iPSCs)*

iPSCs were differentiated to muscle progenitor cells (MPC) using the STEMdiff™ Myogenic Progenitor Supplement Kit (Stem Cell Technologies) for 30 days according to the manufacturer's instructions. Briefly, prior to harvest iPSCs were pre-treated with 10 µM ROCK inhibitor (Y-27632 2HCl, SelleckChem) for 2 hours, washed once with DPBS (1X, without Ca/Mg) and dissociated to single cells using 1X TrypLETM Express (Thermo Fisher). Cells were counted with a haemocytometer and 2.5 – 3.5 x $10^5$ cells/well (line dependent concentration determined empirically) were plated in triplicate into GFR Matrigel-coated wells in a 6-well plate. After 24-hours, evenly plated iPSCs at a suitable density (clusters of ~10-30 cells) were changed into STEMdiff™ Myogenic Progenitor Supplement differentiation media, and media changed daily for 30 days according to the manufacturer's guidelines.

At Day 30, cultures were again pre-treated with 10 µM ROCK inhibitor (SelleckChem) for 2 hours, washed once with DPBS (1X, without Ca/Mg) and treated with 1X TrypLETM Express (Thermo Fisher) for 10 min. Cells were subsequently mechanically dissociated for ~1 min/well using two 25-gauge needles. Collagenase Type IV (2 mg, Cat#17104019, Thermo Fisher) was added to each well using a wide-bore P1000 tip and mixed by pipetting up and down for ~1 min/well, followed by incubation at 37 °C, 5% $CO_2$, for 10 min. Cultures were again mechanically dissociated for ~1 min/well using two 25-gauge needles, followed by pipetting for 1-2 min using a standard P1000 tip. Dissociated cultures were collected into 5 mL of complete MyoTonic™ medium (MyoTonic™ basal medium (Cook Myosite) supplemented with MyoTonic™ serum-free growth supplement (Cook Myosite), 10% FBS (Gibco, Australian Origin), 1% Pen/Strep (in-house) and 20 ng/mL FGF2 (PeproTech, Cat#100-18B)) with 10 µM ROCK inhibitor (Selleckchem, Cat#S1049) and filtered through a 70 µM cell strainer. The resulting single cells were termed myogenic progenitor cells (MPC) and were pelleted at 300g for 5 min at RT with low brake (2), resuspended in complete MyoTonic™ supplemented with 10 µM ROCK inhibitor and counted. Trypan blue was used to assess cell viability (should exceed 80%) and 1.5-2.5 x $10^5$ viable cells/well plated into Matrigel-coated (standard formulation, 1:200 in DMEM/F-12; Thermo Fisher) 6-well plates or cryopreserved as a P0 stock in complete MyoTonic™ supplemented with 20% FBS and 10% DMSO.

*Gymnotic delivery of ASOs to myotubes (extended methods)*

MPCs were cultured and plated for terminal differentiation as described above. For each experiment, cells were plated in triplicate into three different 96-well plate types corresponding to their final downstream analysis requirements; for analysis of gene expression, cells were plated into standard culture plates (Plate 1, Greiner, Cat# 655180); for analysis of cell viability, cells were plated into white culture plates (Plate 2, Greiner, Cat#655083); and, for immunohistochemistry cells were plated on optically clear plates (Plate 3, Thermo Scientific, Cat#165305). On day 0, growth media from all three plates was replaced with freshly prepared myotube differentiation media supplemented with ASOs to a final concentration of 0.25 µM, 0.5 µM or 1.0 µM. ASOs were added to each plate in technical duplicate. Media was not changed, and plates were harvested at D5 of differentiation.

Cell lysate was harvested from Plate 1 and cDNA was generated using the PowerSYBR Green cells-to-Ct kit (Invitrogen, Cat#4402955) according to the manufacturer's instructions. Final cDNA was diluted 1:2 with UltraPure $H_2O$ and stored at -20°C. *MALAT1* transcript knockdown was subsequently analysed by qRT-PCR as described above. All data were first normalised to the untreated condition (within biological replicate), then normalised to the appropriate negative control condition.

Cells in plate 2 were assayed for viability using the CellTiter-Glo® 2.0 Cell Viability Assay (Promega) according to the manufacturer's instructions. Luminescence readings were recorded using the CLARIOstar plate reader (BMG Labtech). The average of four readings per well was calculated, and then technical duplicates were averaged within condition. A media only condition was used to calculate the background luminescence, and this value was subtracted from all other conditions. Data were subsequently normalised to the appropriate negative control condition.

Cells in plate 3 were washed, fixed, stained and imaged on the CellInsight CX7 High Content Analysis Platform (Thermo Scientific) as described above.

**ASOs used in this study**.

| Target | Chemistry | Sequence (5'-3') | Supplier | Catalogue (LOT#) |
|---|---|---|---|---|
| hsMALAT1 | LNA gapmer, PS^, 5'FAM | CGTTAACTAGGCTTTA | QIAGEN | 339516LG00000003-DFB (30702766-1) |
| Neg A | LNA gapmer, PS^ | AACACGTCTATACGC | QIAGEN | 339516LG00000002-DFA (40103151-3) |
| Toxic LNA-1 | LNA gapmer, PS^ | TCGCCGCGGAGGGGAAG | IDT | Custom design |

^PS = Phosophorothioate. All nucleotide linkages are PS modified.

*Mantarray stimulation protocols*

Stimulation and recording of Mantarray data was performed every ~24 h from day 6 to 21. Twitch (1Hz) was always performed first, followed by force-frequency (with a 2-5 min delay between).

**Force-frequency stimulation protocol (total duration 107 sec):**

| Step # | Type | Phase 1 | Phase 2 | Pulse frequency | Active duration |
|---|---|---|---|---|---|
| 1 | Delay (5 sec) | - | - | - | - |
| 2 | Biphasic | 5 ms / 100 mA | 5 ms / -100 mA | 1 Hz | 2 sec |
| 3 | Delay (8 sec) | - | - | - | - |
| 4 | Biphasic | 5 ms / 100 mA | 5 ms / -100 mA | 2 Hz | 2 sec |
| 5 | Delay (8 sec) | - | - | - | - |
| 6 | Biphasic | 5 ms / 100 mA | 5 ms / -100 mA | 3 Hz | 2 sec |
| 7 | Delay (8 sec) | - | - | - | - |
| 8 | Biphasic | 5 ms / 100 mA | 5 ms / -100 mA | 5 Hz | 2 sec |
| 9 | Delay (8 sec) | - | - | - | - |
| 10 | Biphasic | 5 ms / 100 mA | 5 ms / -100 mA | 10 Hz | 2 sec |
| 11 | Delay (8 sec) | - | - | - | - |
| 12 | Biphasic | 5 ms / 100 mA | 5 ms / -100 mA | 20 Hz | 2 sec |
| 13 | Delay (8 sec) | - | - | - | - |
| 14 | Biphasic | 5 ms / 100 mA | 5 ms / -100 mA | 30 Hz | 2 sec |
| 15 | Delay (8 sec) | - | - | - | - |
| 16 | Biphasic | 5 ms / 100 mA | 5 ms / -100 mA | 40 Hz | 2 sec |
| 17 | Delay (8 sec) | - | - | - | - |
| 18 | Biphasic | 5 ms / 100 mA | 5 ms / -100 mA | 50 Hz | 2 sec |
| 19 | Delay (8 sec) | - | - | - | - |
| 20 | Biphasic | 5 ms / 100 mA | 5 ms / -100 mA | 80 Hz | 2 sec |
| 21 | Delay (8 sec) | - | - | - | - |
| 22 | Biphasic | 5 ms / 100 mA | 5 ms / -100 mA | 80 Hz | 2 sec |

**Twitch (1 Hz) stimulation protocol (total duration 136 sec):**

| Step # | Type | Phase 1 | Phase 2 | Pulse frequency | Active duration |
|---|---|---|---|---|---|
| 1 | Delay (4 sec) | - | - | - | - |
| 2 | Biphasic | 5 ms / 100 mA | 5 ms / -100 mA | 1 Hz | 30 sec |
| 3 | Repeat above (Steps 1-2) three more times (total duration: 136 seconds) | | | | |

*EMT harvesting and immunostaining (extended methods)*

On harvest days (D7, D14 and D21), tissues were rinsed in cold 1X DPBS and either a) flash frozen in liquid nitrogen (for RNA/protein work), or b) rinsed in cold 1X DPBS, then fixed in 4% PFA overnight at 4 °C (for immunostaining). Following fixation, tissues were stored in 1X DPBS overnight at 4 °C, then dehydrated in 20% sucrose overnight at 4 °C. EMTs were then blotted dry, embedded in OCT in a Tissue-Tek Biopsy Cryomold, and frozen at -80 °C. Tissues were cryosectioned (30 µm) using a Leica CM1860 cryostat.

For immunostaining, frozen cryosections (on slide) were removed from storage (-80 °C) and left to air-dry at room temperature (~15 min). Sections were washed with 1X PBS for 5 min, then repeated for a total of 3x 5 min washes. PBS was removed thoroughly by tilting and aspiration, then air-drying for 5 min. Sections were blocked in blocking buffer (2% BSA + 0.3% Triton X-100 in PBS) for 1 h at RT in a wet chamber, then blocking buffer removed and replaced with primary antibody (diluted in blocking buffer and incubated overnight at 4 °C in a wet chamber). Following overnight incubation, slides were washed with 1X PBS three times for 5 min at RT. Slides were transferred back to wet chamber and secondary antibody (diluted in blocking buffer) added, then incubated for 1 h at RT in a dark chamber. Slides were washed three times for 10 min in PBS at RT. DAPI (1 µM) was added for 10 min at RT in a dark chamber, then slides washed in PBS a further three times for 5 min. Finally, sections were covered with mounting medium (abcam #ab104135), incubated for 5 min, then edges sealed with nail polish. Images were collected on a Leica SP8 confocal microscope. At least 2 – 3 sections were collected, stained and imaged for each tissue.
